# Dendritic Architecture Enables de Novo Computation of Salient Motion in the Superior Colliculus

**DOI:** 10.1101/2025.02.28.640764

**Authors:** Norma K. Kühn, Chen Li, Natalia Baimacheva, Janne Zimmer, Katja Reinhard, Vincent Bonin, Karl Farrow

**Affiliations:** VIB-Neuro-Electronics Research Flanders, Leuven, Belgium; Department of Biology and Leuven Brain Institute, KU Leuven, Leuven, Belgium; Yale School of Medicine, New Haven, CT, USA; Scuola Internazionale Superiore di Studi Avanzati, Trieste, Italy; imec, Leuven, Belgium

## Abstract

Dendritic architecture plays a crucial role in shaping how neurons extract behaviorally relevant information from sensory inputs. Wide-field neurons in the superior colliculus integrate visual information from the retina to encode cues critical for visually guided orienting behaviors. However, the principles governing how these neurons filter their inputs to generate appropriate responses remain unclear. Using viral tracing, two-photon calcium imaging, and computational modeling, we show that wide-field neurons receive functionally diverse inputs from twelve retinal ganglion cell types, forming a layered, type-specific organization along their dendrites. This structured arrangement allows wide-field neurons to multiplex salient motion cues, selectively amplifying movement and suppressing static features. Computational models reveal that the spatial organization of dendrites and inputs enables the selective extraction of behaviorally relevant stimuli, including *de novo* computations. Our findings underscore the critical role of dendritic architecture in shaping sensory processing and neural circuit function.

## Introduction

Dendrites add a spatial component to the computations a neuron can perform, expanding its processing capabilities beyond temporal integration at the cell body. Dendritic inputs are often highly structured, with laminar arrangements aligning branches with inputs from specific brain regions or cell types^1–4^ and sensory areas like the visual cortex and superior colliculus exhibiting retinotopic organization^5–8^. These spatial arrangements dictate where information enters the dendritic tree and how it is processed through passive and active electrical properties, enabling a set of linear and nonlinear computations, e.g., band pass filtering^9,10^ and nonlinear summation^11–14^. Studies in local circuits of the retina have linked dendritic architecture and input location to specific computations of defined cell types^15–17^. However, our understanding of how these factors shape computations as information is passed on to central brain regions remains poor, largely due to limited knowledge of the relationship between specific inputs^1,18^ and dendritic structure^19,20^. Here, we demonstrate that dendritic architecture and arrangement of inputs enable a *de novo* computation of salient motion cues in neurons of the mouse superior colliculus (colliculus), a brain area that links identified retinal inputs with molecularly and morphologically defined cell types.

In the colliculus, saliency is strongly tied to object motion^21–23^, enabling mice to rapidly detect approaching threats or fleeing prey^24,25^. Neurons in the colliculus integrate inputs from approximately 40 retinal ganglion cell types^26–28^, each encoding distinct visual attributes, motion, contrast, or orientation, to guide innate orienting behaviors, including predator avoidance and prey capture^24,25,29,30^. A key question is whether the colliculus merely relays visual features coming from the retina or computes new ones. The retinotopic organization of the superior colliculus ensures that these inputs are spatially preserved, creating a detailed map of the visual field^7,31,32^. Additionally, the specific depth at which retinal inputs terminate in the optic layers is hard-wired^2,31,33,34^, which could enable collicular neurons to sample specific inputs across depth. Recent evidence suggests that the colliculus does not merely inherit visual representations from the retina but selectively filters specific features to generate saliency maps^31,35–38^.

Here, we investigate how genetically-targetable wide-field neurons of the colliculus process and refine their retinal inputs to extract behaviorally relevant information. Wide-field neurons extract motion saliency^39,40^ from a defined set of retinal ganglion cell types^41^ to guide innate freezing and hunting behaviors (Figure 1A)^25,42–44^. To identify how dendrites shape the retinal inputs, we mapped the direct inputs and outputs of wide-field neurons in response to a set of visual stimuli, using viral tracing, two-photon calcium imaging and patch-clamp recordings. We found that the unique dendritic structure of wide-field neurons, along with the spatial distribution of their inputs enables multiplexing of salient motion cues critical for detecting both predators and prey. While some retinal ganglion cells exhibit strong selectivity for looming stimuli^45^, signaling potential predators, the representation of receding motion, essential for tracking escaping prey, is weak in the retinal signals and emerges *de novo* within the colliculus. Computational modeling revealed that, contrary to expectations, nonlinear processing beyond a basic rectifier plays a minimal role. Instead, the summation of retinotopically organized inputs across the dendritic tree is key to these computations. Our findings highlight the spatial organization of dendrites and their inputs as a fundamental substrate for sensory processing, supporting the colliculus’s role in computing and propagating motion saliency essential for survival.

**Figure 1.**
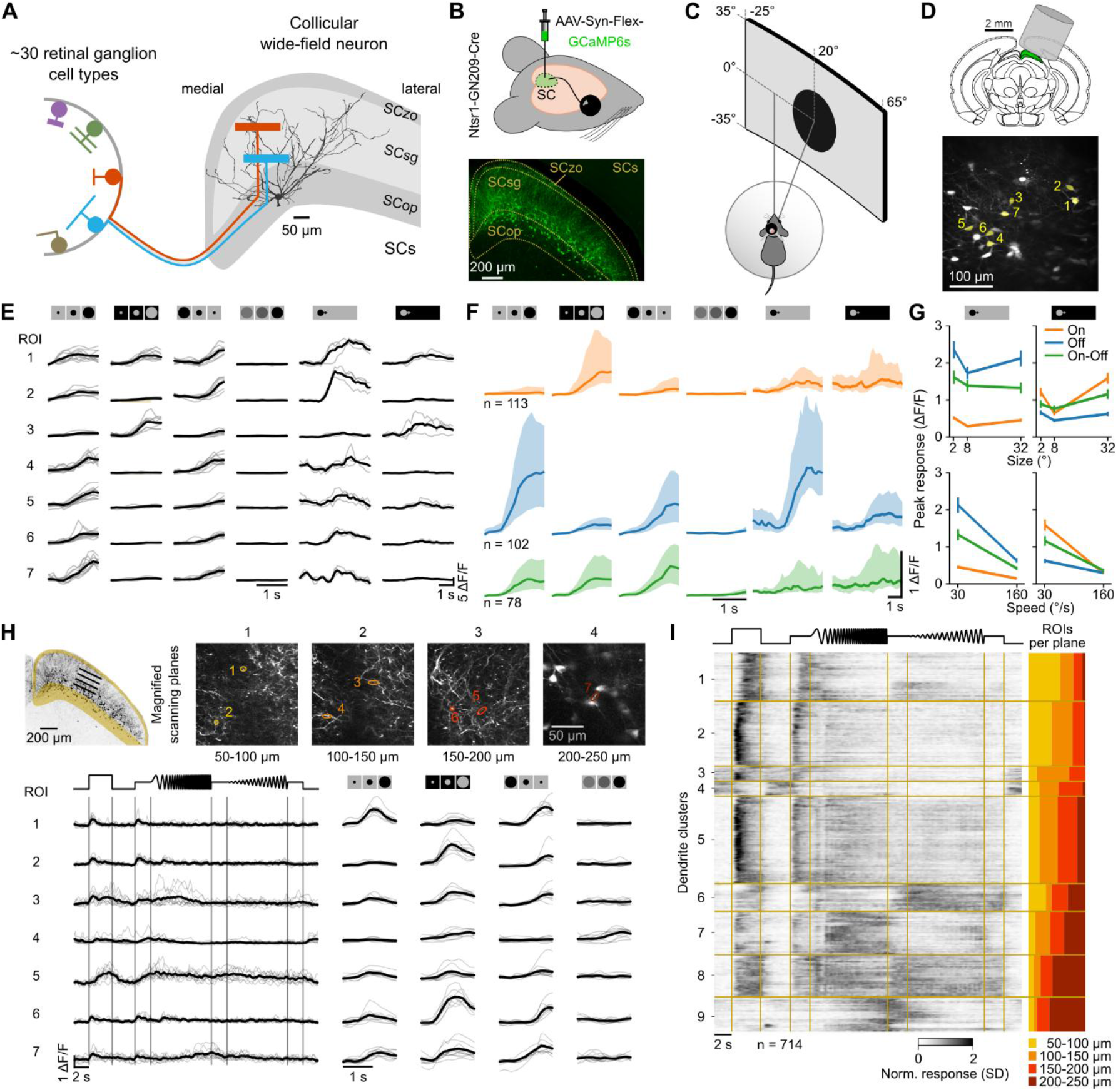
Characterization of Visual Responses of Wide-Field Cell Bodies and Dendrites. (A) Schematic of wide-field neurons of the superior colliculus and type-specific layered input from retinal ganglion cells. (B) Viral labeling of wide-field neurons in Ntsr1-GN209-Cre mice. SCzo, zonal layer; SCsg, superficial gray layer; SCop, optic layer of the sensory-related superior colliculus (SCs). (C) In vivo two-photon calcium imaging setup. Visual stimuli were presented at 0° elevation and 20° azimuth in front of the contralateral eye. (D) Position of the cranial window for two-photon imaging (top) and example field of view (bottom). (E) Wide-field neuron cell body responses; individual trials shown in gray, median response in black. Cell body locations indicated in (D). Visual stimuli from left to right: black and gray expanding disk, black shrinking disk, dimming disk, and black and gray sweeping disks. (F) Median and interquartile range of clustered On, Off, and On-Off subtype responses to expanding, shrinking, dimming, and sweeping disks. (G) Mean peak responses of On, Off, and On-Off types to dark and bright sweeping stimuli. Data from 12 mice (293 cell bodies). (H) Scanning planes for volumetric imaging (top) and example dendritic ROI responses across all scanning planes (bottom). Scanning planes magnified ×2; ellipses indicate ROIs. (I) Heatmap of normalized and clustered dendritic responses to a full-field stimulus (left) and depth profiles of identified clusters (right), sorted by mean cluster depth. Data from 18 mice across 22 sessions (714 ROIs). See also Figure S1–S3.

## Results

### Wide-Field Neurons Multiplex Motion Signals and Exhibit Diverse Contrast Preferences

We investigated the role of dendrites in wide-field neuron encoding of salient stimuli in three steps: (1) recording the responses of wide-field neurons and dendritic compartments, (2) identifying the visual features relayed by innervating retinal ganglion cells, and (3) determining the key components required to transform inputs into outputs using computational models incorporating dendritic properties. We assessed the visual features encoded by wide-field neurons by recording their responses to visual stimuli in head-fixed, behaving mice using two-photon calcium imaging (Figure 1). A calcium indicator was expressed using a Cre-dependent adeno-associated virus (AAV-Syn-Flex-GCaMP6s) injected into the left colliculus of *Ntsr1-GN209 Cre* mice (Figure 1B). Visual stimuli were presented to the contralateral eye, while the mouse was free to run on a floating ball (Figure 1C). Neurons were imaged through a cannula window, implanted above the anterior two-thirds of the colliculus (Figure 1D).

We found that wide-field neurons respond to a diverse set of moving stimuli but showed minimal responses to static stimuli (Figure 1E). To characterize their response properties, we presented expanding, shrinking and sweeping black disks, as well as dimming disks on a gray background (see *Methods & Materials*). Disks were linearly expanding or shrinking with an edge speed of 30°/s, while sweeping disks were presented in three sizes (2°, 8°, 30°) and two speeds (30°/s and 160°/s). Expanding and sweeping disks were also shown in reverse contrast. Calcium responses were recorded from wide-field neuron cell bodies at 200-250 μm below the surface of the colliculus (Figure 1D-E). Wide-field neurons exhibited diverse contrast preferences, with some responding only to black moving disks on a gray background and others only to gray disks on black (Figure 1E). Their responses were strongest to slowly moving edges, as indicated by comparable peak responses to expanding, shrinking and slowly sweeping disks. Peak responses to gray expanding, black shrinking, dimming and black and gray slowly sweeping disks, computed from responses normalized by the standard deviation (SD) across stimuli, were compared to those elicited by black expanding disks (median [interquartile range, IQR] difference in SD: −0.05 [−0.27, 0.18], −0.08 [−0.26, 0.07], 0.11 [−0.13, 0.31], −0.15 [−0.36, 0.04]; p values 0.48, 0.28, 0.14, 0.34, Kruskal-Wallis test). In contrast, responses were significantly weaker for dimming disks and fast sweeping disks (median [IQR] difference: −0.61 [−0.76, −0.49], −0.43 [−0.60, −0.28], −0.44 [−0.61, −0.31]; p values 10^−38^, 10^−10^, 10^−9^, Figure S1). These results show that selective encoding of slowly moving edges enables wide-field neurons to act as multiplexers of motion cues.

We identified three functional types: On, Off, and On-Off cells, based on clustering and contrast preference to expanding, shrinking, and dimming disks (Figure 1F-G, Figure 1, see Methods & Materials). Each type responded strongly to expanding, shrinking and sweeping disks of their preferred contrast but not to the dimming disk where no moving edge was present (Figure 1F-G). On and Off types showed significantly stronger peak responses to an expanding disk of their preferred contrast compared to an expanding disk of their unpreferred contrast (median [IQR] difference of peaks: On: 1.7 [1.4, 1.9] SD, Off: 1.3 [1.0, 1.6] SD, p values 10^−21^ and 10^−25^, Kruskal-Wallis test), while On-Off cells did not show statistically significant differences between On and Off contrasts (p = 0.09). All wide-field neuron types showed a preference for slower speeds of the sweeping stimuli, as observed in higher peak responses for a speed of 30°/s compared to a speed of 160°/s for most stimuli (median [IQR] differences of peaks between slow and fast: 0.55 [0.41, 0.72], 0.23 [0.14, 0.35], 0.32 [0.25, 0.39] for black and 0.09 [0.06, 0.16], 0.01 [−0.02, 0.05], 0.22 [0.15, 0.43] gray disks of sizes 2°, 8° and 32°, p values 10^−18^, 10^−9^, 10^−14^, 0.06, 0.42, 10^−12^, Wilcoxon signed rank; Figure 1G). The preference for slow speeds of sweeping disks is in line with the ratio of wide-field neurons’ preferred spatial and temporal frequencies that peak at 2 Hz and 0.04/°, respectively, resulting in a preferred speed of ~30°/s (Figure S2). This suggests that wide-field neurons are finely tuned to process slow motion signals critical for detecting environmental changes, such as the movement of predators or prey, with their selective encoding emphasizing the saliency of slow motion cues.

### Depth-Dependent Clustering of Dendritic Visual Responses

Retinal inputs to the colliculus are organized in distinct functional and depth-specific clusters^2,33^. A key question is whether wide-field neuron dendrites maintain the layered organization of retinal inputs, potentially enabling compartmentalized processing of visual information. To investigate this, we presented a full-field stimulus with rich temporal dynamics but no moving edges^18,26^ and recorded dendritic calcium responses quasi-simultaneously across four scanning planes, from the collicular surface to the cell bodies (Figure 1H). The stimulus evoked highly localized responses in dendrites (Figure 1H, see also Figure S3).

Clustering dendritic responses based on correlation analysis revealed nine distinct functional groups (Figure 1I). Seven clusters exhibited On-type responses to the transition from black to white, while only cluster 3 and 4 showed Off-type responses with sustained activity when the screen turned black. The clusters also differed in temporal frequency preference, with most responding to low frequencies (clusters 1–6), whereas clusters 7 and 8 were tuned to higher frequencies.

These functional clusters were organized by depth, suggesting a functional compartmentalization of the dendrites. Sorting clusters by the mean depth of their signals revealed a characteristic depth profile (Figure 1I, right, see Methods & Materials): clusters 1 and 2 occupied the most superficial layers (50–150 μm), clusters 3 and 4 were found at intermediate depths (100– 200 μm), clusters 5–7 spanned all scanning planes, and clusters 8 and 9 were restricted to the deepest layers (150–250 μm). This structured depth-dependent organization suggests that different retinal ganglion cell types provide depth-specific input to wide-field neuron dendrites, potentially allowing distinct visual features to be processed in separate dendritic compartments.

### Twelve Retinal Ganglion Cell Types Relay Diverse Visual Features to Wide-Field Neurons

To identify the retinal ganglion cells that provide input to wide-field neurons, we used rabies-based transsynaptic retrograde tracing to label innervating retinal ganglion cells (Figure 2A, see also^41^). Labeled cells were recorded *ex vivo* using two-photon calcium imaging or whole-cell patch-clamp recordings while visual stimuli, identical to those presented to wide-field neurons, were projected onto the retina (Figure 2B-C). To ensure consistency in the dataset and facilitate comparison with calcium signals in wide-field neurons, spiking activity from patch-clamp recordings was converted to calcium signals by convolving with a GCaMP6s kernel (rise time: 0.1 s, decay time: 0.4 s; see *Methods & Materials*).

**Figure 2.**
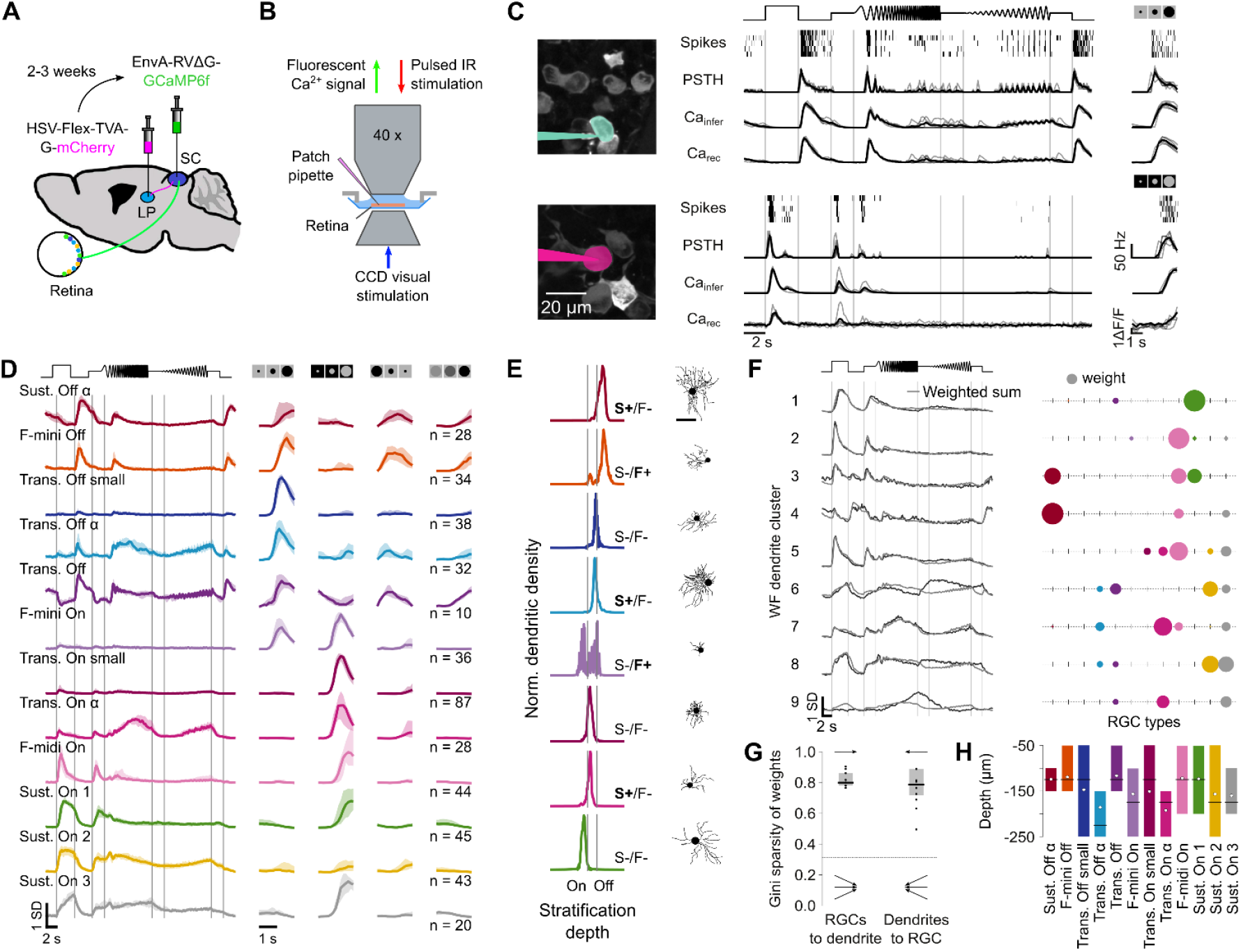
Twelve Retinal Ganglion Cell-Types Provide Input to Wide-Field Neurons. (A) Viral strategy to label retinal ganglion cells innervating wide-field neurons. Retrograde helper virus (HSV-Flex-TVA-G-mCherry, magenta) was injected into the lateral posterior nucleus of the thalamus (LP) and rabies virus (EnvA-RVΔG-GCaMP6f, green) into the superior colliculus (SC). (B) Ex vivo recordings of labeled retinal ganglion cells using two-photon calcium imaging and whole-cell patch-clamp recordings. (C) Spiking responses, peristimulus time histogram (PSTH), inferred calcium signal and simultaneously recorded calcium signals of a sustained Off alpha cell (top) and a transient On alpha cell (bottom). (D) Normalized responses of identified retinal ganglion cell clusters (lines, median; shaded areas, interquartile range). Data from 24 mice (445 retinal ganglion cells) (E) Examples of morphologically and molecularly identified retinal ganglion cell types, color-matched to clusters in (D). S+/−, SMI32 positive or negative; F+/−, FOXP2 positive or negative. (F) Weighted sum of retinal ganglion cell types to match full-field stimulus responses of wide-field neuron dendrites. Left: Weighted sum (gray) to match median response of each dendritic cluster (black). Right: Weights of each retinal ganglion cell type. Disc color and size indicate cell type and weight, respectively. (G) Gini sparsity of weights of retinal ganglion cell types to wide-field dendrite or wide-field dendrites to retinal ganglion cell type. Horizontal line indicates Gini sparsity of uniformly distributed weights. (H) Depth distribution of retinal ganglion cell inputs in the colliculus, determined by the most strongly correlated dendritic signals of wide-field neurons. Gray dots, horizontal lines and boxes indicate mean, median and interquartile range of each ganglion cell type. See also Figure S4 and Methods & Materials.

Examples of simultaneously recorded spiking and calcium activity, along with convolved signals, are shown in Figure 2C (see also Figure S4). To classify the recorded retinal ganglion cells based on molecular and structural features, a subset of cells was stained for key molecular markers of major retinal ganglion cell types (Smi32, Foxp2, Cart), and their dendritic arbors were reconstructed (see *Methods & Materials*, Figure S4).

Retinal ganglion cell calcium responses were grouped using agglomerative clustering with subsequent manual curation (Figure 2D, see *Methods & Materials*), leading to twelve functional types, five Off, one On-Off and six On types (Figure 2D). Clusters were matched to existing cell types from previous studies^26,27,41^ by using a subset of molecularly and morphologically identified cells (Figure 2E, see also Figure S4).

The identified On, Off, and On-Off retinal ganglion cell types exhibit distinct response profiles to full-field flashes and expanding disks of corresponding contrast. On types respond significantly more strongly to On flashes and expanding gray disks compared to Off flashes and expanding black disks, while Off types show the reverse pattern. On-Off types respond to both black and gray expanding disks, indicating sensitivity to both contrast polarities. In addition to their responses to expanding black disks, four of the five Off cell types also responded to shrinking and dimming disks, though with lower amplitudes (median [IQR] differences of peaks −0.32 [−0.4, −0.2] and −0.57 [−0.67, −0.47] SD, p values 0.002 and 10^−21^, Kruskal-Wallis test). This response profile contrasts with that of wide-field neuron cell bodies, where all three subtypes showed strong responses to shrinking black disks but did not respond to dimming disks (Figure 1F). These differences suggest that wide-field neurons integrate retinal inputs in a manner that selectively amplifies specific motion cues while filtering out luminance changes.

### Modeling and Correlation Analysis Reveal Layered, Type-Specific Input to Wide-Field Neuron Dendrites

To determine whether the diverse response profiles observed in wide-field neuron dendrites can be explained by the retinal inputs, we fitted a linear regression model with non-negative matrix factorization (NNMF, see *Methods & Materials*), to the full-field stimulus responses of each wide-field dendrite cluster as a weighted sum of the twelve retinal ganglion cell types. The positively constrained weights reflect the excitatory nature of retinal inputs. This approach produced reasonable fits and a sparse weight matrix (Figure 2F-G). Responses in the upper dendrites were well reconstructed by the linear model (Figure 2F, cluster 1-5 and 7), whereas responses in the lower dendrites showed poorer fits (cluster 6, 8 and 9). This suggests that summation closer to the cell body may involve additional integration mechanisms (Figure 2F).

The fitted weights reveal that each dendritic signal can be accounted for by only a few retinal ganglion cell types (Figure 2F). To quantify the convergence and divergence of retinal inputs onto wide-field dendrites, we calculated the Gini sparsity index (Figure 2G, see *Methods & Materials*). High values (0.81 [0.78, 0.86] for retinal ganglion cell-to-dendrite connections and 0.77 [0.72, 0.83] for dendrite-to-retinal ganglion cell connections) indicate sparse, highly specific connectivity, significantly deviating from a uniform distribution (p values 10^−38^ and 10^−21^, respectively, Mann-Whitney U test, Figure 2G). This further supports functional compartmentalization, where each local dendritic signal is shaped by only a few retinal ganglion cell types. Correlating retinal signals with signals measured in the dendrites of wide-field neurons with the most strongly correlated dendritic signals determining their depth, revealed that retinal ganglion cell types map onto specific dendritic locations within the colliculus (Figure 2H). These findings demonstrate that retinal inputs are organized in a layered, type-specific manner along wide-field neuron dendrites, supporting functional compartmentalization.

### A Simple Weighted Sum of Retinal Inputs Fails to Account for Wide-Field Neuron Peak Responses

Wide-field neurons perform a *de novo* computation of responses to shrinking disks that cannot be explained by a simple weighted sum of retinal inputs. Unlike dendritic signals, cell body responses cannot be reconstructed using such a sum. To test whether they fit a linear model with non-negative weights, we reconstructed responses to both contrasts of expanding disks, as well as shrinking and dimming disks (Figure 3A, see Methods & Materials). The fitted weights indicate that each wide-field neuron subtype receives input from a limited set of retinal ganglion cell types. On and Off subtypes appear to be driven by largely distinct sets of retinal inputs with opposite weight distributions, while the On-Off subtype shares inputs with both On and Off subtypes (Figure 3B, see also Figure S5). Notably, while the On subtype’s responses were well-matched by the linear model, the model failed to capture Off and On-Off subtype responses to a shrinking disk (Figure 3C).

**Figure 3.**
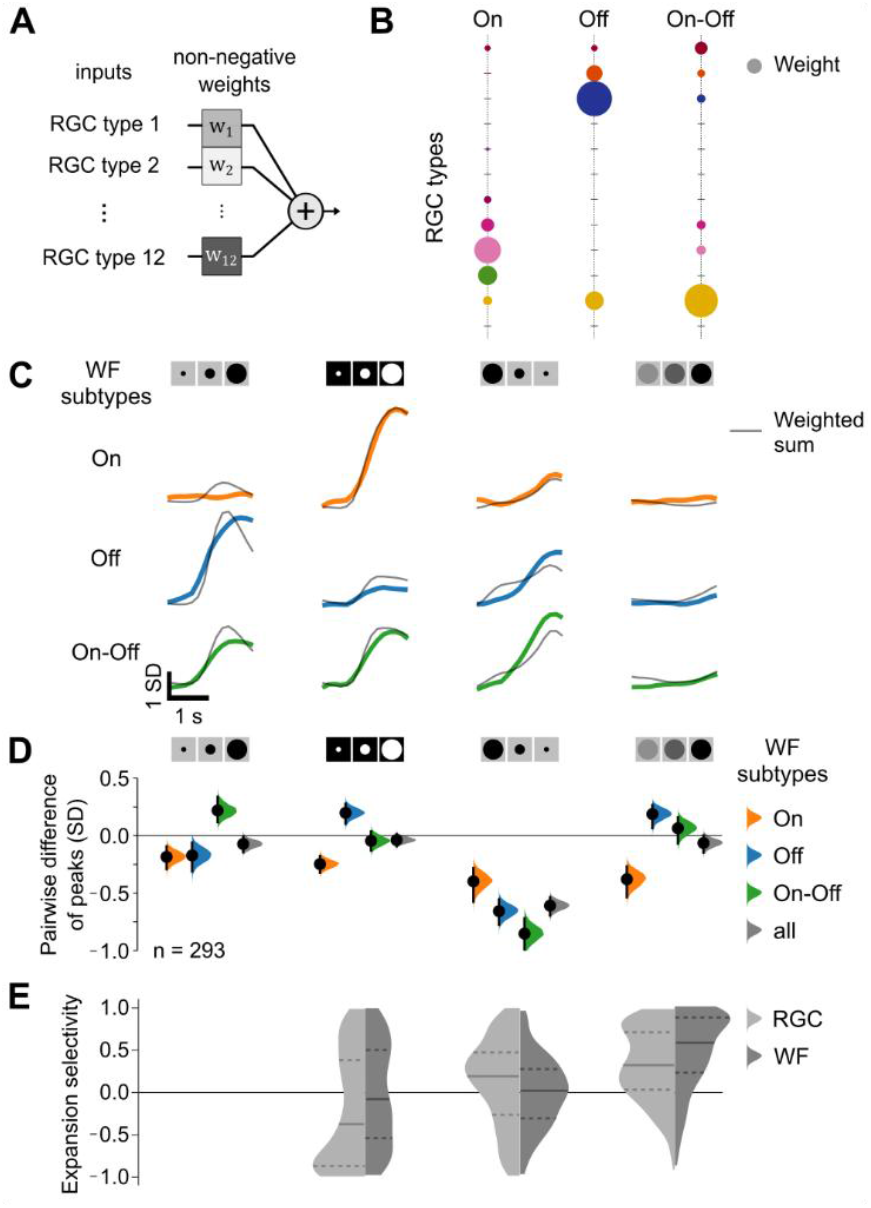
A Linear Point Neuron Model Underestimates Wide-Field Neuron Responses to Shrinking Stimuli. (A) Non-negative linear regression model predicting wide-field neuron cell body responses from retinal ganglion cell inputs. (B) Fitted weights for each retinal ganglion cell type, indicating their contribution to wide-field neuron subtypes. (C) Mean linear model fits (gray) overlaid on measured responses (colored) for the three wide-field neuron subtypes. (D) Peak response differences between measured and fitted responses for each wide-field neuron subtype. Dots and vertical lines indicate the mean and 95% confidence interval of the resampled distribution. (E) Expansion selectivity index distributions for retinal ganglion cells (light gray) and wide-field neurons (dark gray). An index of 1 indicates a strong preference for dark expanding disks. Horizontal and dashed lines represent the median and interquartile range, respectively. Data as in Figures 1 and 2. See also Figure S5.

To quantify the discrepancy between peak amplitudes of measured and fitted responses, we calculated their difference for each wide-field neuron subtype. The distribution of these differences revealed a systematic underestimation of peak responses to the shrinking stimulus especially in Off and On-Off subtypes. A similar trend was observed when calculating the mean squared error for each stimulus type (Figure 3D, see also Figure S5).

Since the response amplitude of wide-field neurons to shrinking stimuli is consistently underestimated by the weighted sum of retinal inputs, we asked whether the strong responses reflect a *de novo* computation within wide-field neurons. Retinal ganglion cells showed stronger responses to expanding stimuli, suggesting that the enhanced shrinking responses in wide-field neurons may emerge within the colliculus. To test this, we computed an expansion selectivity index based on peak responses for each retinal ganglion cell and wide-field neuron and compared their distributions (Figure 3E, see *Methods & Materials*). This revealed a shift in stimulus preference: while retinal ganglion cells preferred expanding over shrinking stimuli (expansion selectivity index: 0.17 [−0.26, 0.46]), wide-field neurons responded similarly to both (expansion selectivity index: 0.02 [−0.26, 0.27]) with significant difference from the retinal distribution (p = 10^−4^, Mann Whitney U). Furthermore, wide-field neurons exhibited a stronger preference for expanding over dimming stimuli (0.54 [0.17, 0.83]) compared to retinal ganglion cells (0.31 [0.01, 0.70], p = 10^−6^, Mann Whitney U).

### Accounting for Inhibition from Collicular Interneurons Does Not Improve Predictions of Wide-Field Neuron Responses

Accounting for inhibition that wide-field neurons receive from GABAergic interneurons of the colliculus^29^ is not sufficient to explain the enhanced responses to shrinking stimuli. To test whether local inhibition modulates wide-field neuron responses, we targeted *Gad2*-positive neurons of the colliculus by expressing GCaMP6s in *Gad2-IRES-Cre* mice (Figure 4A). We recorded responses of *Gad2* neurons to full-field, expanding, shrinking, and dimming stimuli and clustered them using the same approach as for wide-field neurons. This revealed four distinct response types (Figure 4B, see also Figure S6). Types 1 and 2 responded most strongly to the full-field stimulus, whereas types 3 and 4 were broadly activated by all stimuli, with a slight preference for On type stimuli, e.g., the bright expanding disk and the On component of the full-field stimulus.

**Figure 4.**
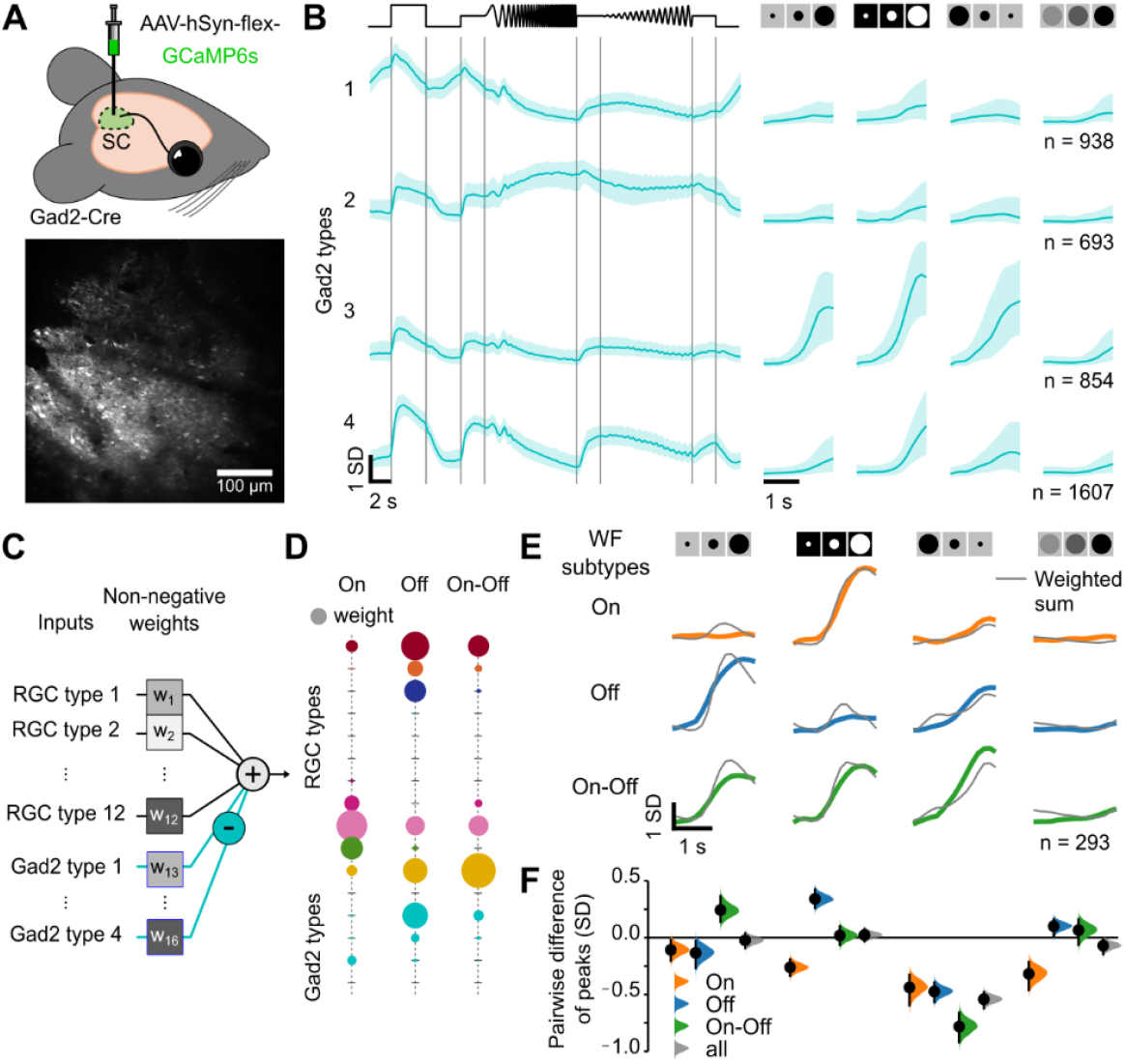
Accounting for Inhibition from GABAergic Neurons in the Colliculus Provides Only Marginal Improvement to the Linear Model. (A) Labeling of Gad2-positive neurons in the colliculus (SC) and an example field of view from in vivo two-photon imaging. (B) Median and interquartile range of normalized responses from four Gad2 neuron clusters to full-field, expanding, shrinking and dimming stimuli. Data from 4096 neurons across 22 sessions from 12 animals. (C) Extended linear model incorporating inhibition, with responses of Gad2 clusters sign-inverted to reflect their inhibitory effect. (D) Fitted weights of retinal ganglion cell and Gad2 neuron types, indicating their contributions to wide-field neuron subtype responses. (E) Weighted sum (gray) overlaid on measured wide-field neuron responses (colored). (F) Pairwise difference of peaks between fitted and measured responses. Dots and vertical lines indicate the mean and 95% confidence interval of the resampled distribution. See also Figure S6.

To incorporate inhibition into our model, we sign-inverted the *Gad2* neuron responses while maintaining non-negative weights, as in the previous model (Figure 4C). This revealed that inhibition primarily originated from *Gad2* types 1 and 2, which contributed to both somatic and dendritic signals of wide-field neurons (Figure 4D, see also Figure S6). In particular, reconstructions of Off subtype wide-field neurons included inhibitory input from type 1 and a smaller contribution from type 2. When fitting the responses of wide-field neuron dendrites, type 1 inhibition contributed more prominently to signals in lower dendritic regions, while type 2 influenced signals in upper dendritic regions (Figure S6).

Even with inhibitory inputs included, the model still underestimated wide-field neuron responses to shrinking stimuli (Figure 4E). While inhibition may shape dendritic processing, it alone does not account for the strong somatic responses to shrinking stimuli.

### Complex Multilayer Models with Nonlinear Integration Offer Only Marginal Improvements

While increasing model complexity enhances fit quality, nonlinear multilayer models provide only limited gains in predicting wide-field neuron responses. To assess the role of nonlinear processing, we implemented a fully connected multilayer perceptron (MLP) that incorporates excitatory and inhibitory summation, as well as nonlinear dendritic and somatic processing. A sigmoidal nonlinearity was applied to model supra- and sublinear integration along the dendrites, while the spiking threshold at the soma was captured using a rectifying nonlinearity at the output layer (Figure 5A, see also Figure S7).

**Figure 5.**
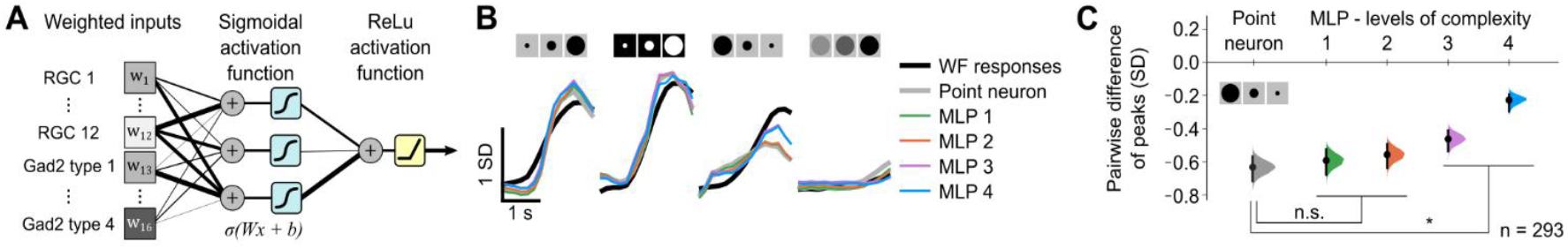
A Multilayer Model with Nonlinearities Improves Fit at the Cost of Complexity. (A) Multilayer perceptron (MLP) model incorporating retinal and inhibitory collicular inputs, with sigmoidal activation functions in hidden layers and a rectifying nonlinearity at the output layer. (B) Mean wide-field cell body responses (colored), with linear fits (black) and MLP fits of increasing complexity: 1: single-layer model with a rectifying output, 2: two-layer model with three hidden sigmoidal nodes, 3: two-layer model with five hidden sigmoidal nodes, 4: three-layer model with five and two hidden nodes in the first and second layers, respectively. (C) Model performance comparison based on the pairwise difference of peaks of wide-field neuron responses and fits for the shrinking stimulus, across models described in (B). Mean and confidence intervals derived from 10,000 bootstrap resamples.

We fitted models of varying complexity, from one to three layers, using least-squares fitting with L1 regularization (see Methods & Materials). The simplest model, which only included a rectifying nonlinearity, showed negligible improvement, whereas additional layers progressively enhanced fit accuracy when considering the average response (Figure 5B). To specifically evaluate model performance on the shrinking stimulus, we analyzed the difference of peaks between observed and predicted responses (Figure 5C). Increasing model complexity gradually reduced this difference, with significant improvements in 2-layer models containing five hidden nodes and 3-layer models with five plus two hidden nodes. However, these models required a high number of parameters and dense weight matrices, performing best only when regularization was minimal (see also Figure S7). This suggests that additional, potentially linear mechanisms must be considered to fully explain wide-field neuron responses.

### Passive Dendritic Properties Improve Predictions of Wide-Field Neuron Responses

Incorporating dendritic filtering and spatial morphology provides a simple yet effective refinement for modeling wide-field neuron responses and can account for *de novo* computation. Passive dendrites act as low-pass filters, introducing temporal delays for distal inputs relative to those near the soma (Figure 6A). Additionally, the spatial arrangement of dendrites and the retinotopic organization of inputs create a spatiotemporal activation pattern, wherein different dendritic regions are sequentially activated during stimulus motion (Figure 6B). To account for these effects, we extended the model by incorporating filtered input channels, effectively doubling the parameter set.

**Figure 6.**
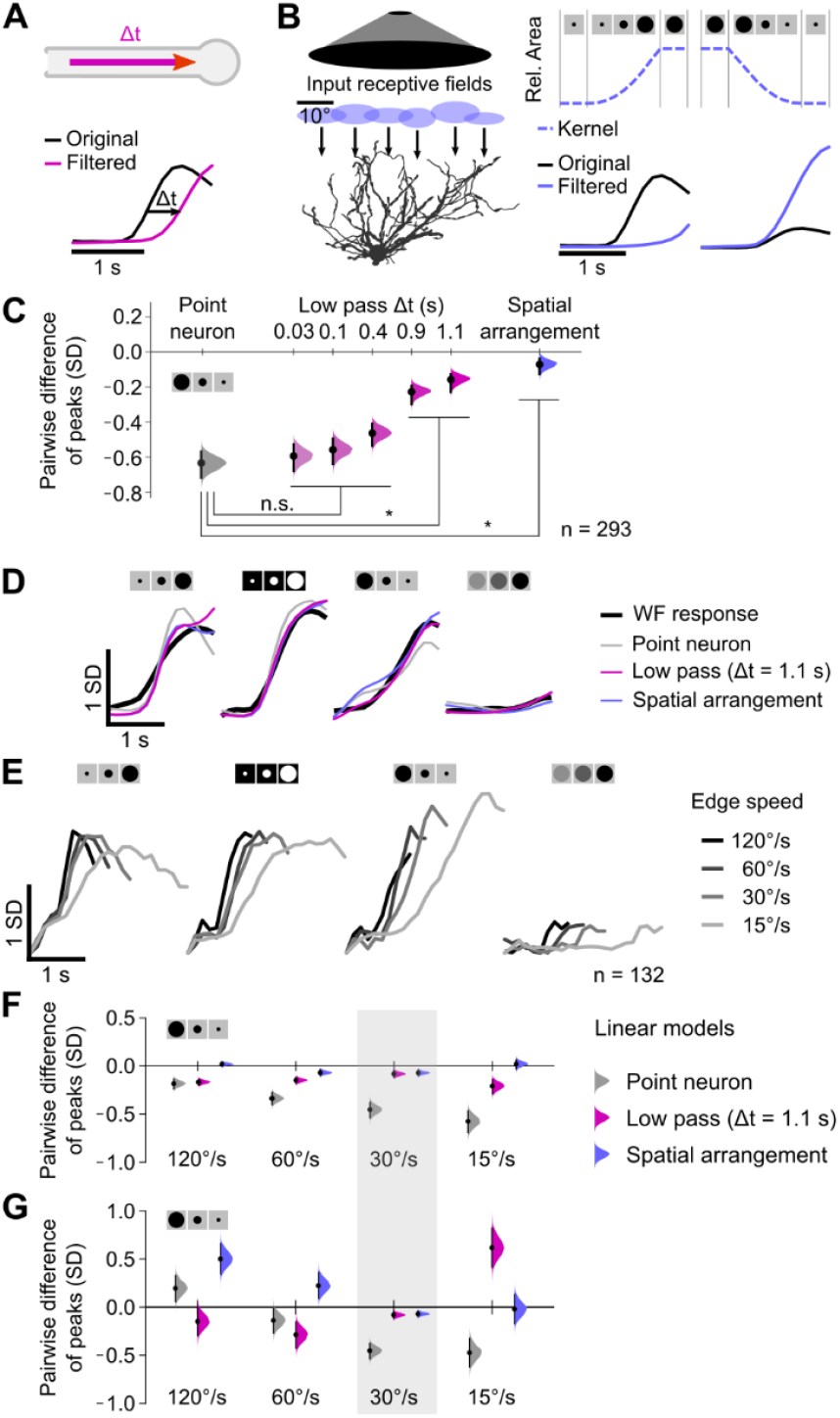
Accounting for Low Pass Filtering and Spatiotemporal Integration of Inputs by Dendrites Improves Model Performance. (A) Low pass filtering of retinal inputs induces signal delay. Original input: black; filtered input: pink. (B) Left: Spatial arrangement of inputs and dendrites. Right: Spatiotemporal activation is modeled by a kernel. Original input: black; filtered input: purple. (C) Pairwise differences between peak responses of wide-field neurons and model fits to the shrinking stimulus. Point neuron: gray; low-pass filtering: pink; spatially arranged dendrites: purple. Black dots with error bars represent mean and confidence intervals from bootstrapping. (D) Mean responses of wide-field neurons (black) and a subset of the linear fits shown in (C). Data as in Figure 1. (E) Mean wide-field neuron responses to expanding, shrinking, and dimming disks at four different edge speeds (15, 30, 60, 120°/s, grayscale-coded). Gray-shaded areas indicate time of movement. (F) Pairwise differences of peaks between wide-field neuron responses and model fits to different speeds. (G) Pairwise differences of peaks as in (F), but with a model fitted to the original speed of 30°/s.

To examine the impact of dendritic filtering, we applied low-pass filters with five different cutoff frequencies (3, 2, 1, 0.5 and 0.4 Hz, see *Methods & Materials*). Lowering the cutoff frequency progressively improved model fits (Figure 6C). Temporal shifts in input signals increased accordingly, with a 0.5 Hz cutoff (0.9 s delay) significantly improving peak response predictions for shrinking stimuli compared to the point neuron model (p values 10^−11^, 10^−6^, 0.58, 0.98, 0.97 of cutoff frequencies 0.4, 0.5, 1, 2 and 3 Hz, Mann-Whitney U test; Figure 6C). The best performance was achieved at 0.4 Hz (1.1 s delay; Figure 6C-D).

To model the spatiotemporal activation of retinotopically distributed inputs, we convolved retinal inputs with a quadratic kernel (Figure 6B; see *Methods & Materials*). This kernel simulates the gradual recruitment of inputs during stimulus expansion or shrinkage, while dimming responses remained unaltered due to constant input recruitment. Despite its simplicity, this phenomenological model accurately reproduced wide-field neuron responses, performing significantly better to the point neuron model and comparable to the best low pass filter model (p values 10^−13^ and, respectively; Figure 6C-D).

To assess the generalizability of our models, we tested them on a withheld dataset featuring expanding, shrinking, and dimming disks presented at three additional edge speeds (15, 30, 60, 120°/s, corresponding to movement durations of 1.9, 0.95, 0.48 and 0.24 s, respectively; see *Methods & Materials*). The original dataset had an edge speed of 30°/s. Across wide-field neurons, the slowest shrinking disk (15°/s) elicited the highest peak amplitudes (Figure 6E). These responses were best captured by the model incorporating the spatial arrangement of inputs and dendrites with peak amplitudes indistinguishable from measured responses (mean difference of peaks 0.02±0.05 SD, p = 0.14, Wilcoxon signed-rank; Figure 6F). This model also provided the most accurate predictions for lower peak amplitudes in response to fast shrinking disks (120°/s), outperforming both the linear point neuron and low pass filter models (mean difference of peaks 0.02±0.03 SD, p = 0.38, Wilcoxon signed-rank; mean difference from point neuron and best low pass filter model: −0.14±0.06 and −0.13±0.06 SD, p values 10^−4^ and 10^−4^, Mann-Whitney U; Figure 6F).

The most effective way to capture wide-field neuron response properties is by accounting for the spatial arrangement of synaptic inputs along their dendrites. Applying a spatiotemporal activation kernel accurately reconstructs peak responses to behaviorally relevant visual stimuli, emphasizing the evolutionary advantage of wide-field neurons’ extensive dendritic arbors.

To further refine our model, we incorporated depth-dependent filtering to account for layer-specific inputs along the dendritic tree (Figure 2H). Retinal inputs to the upper dendritic layers (depth < 150 µm) underwent spatiotemporal filtering, while inputs to the lower layers (> 150 µm) remained unaltered (Figure 7A). This reduced model complexity to just twelve parameters while preserving predictive power, with peak amplitudes indistinguishable from measured responses for almost all speeds (mean peak differences −0.07±0.10, −0.05±0.07, 0.01±0.06 and 0.12±0.04, p values 0.48, 0.36, 0.99 and 10^−5^ for 15, 30, 60 and 120°/s, Wilcoxon signed-rank; Figure 7B), outperforming the point neuron model despite fitting the same number of parameters. By restricting each input to a specific layer, this model minimizes wiring costs while maximizing coding efficiency, underscoring the evolutionary significance of retinotopically organized, layer-constrained inputs in integrating and encoding dynamic visual information.

**Figure 7.**
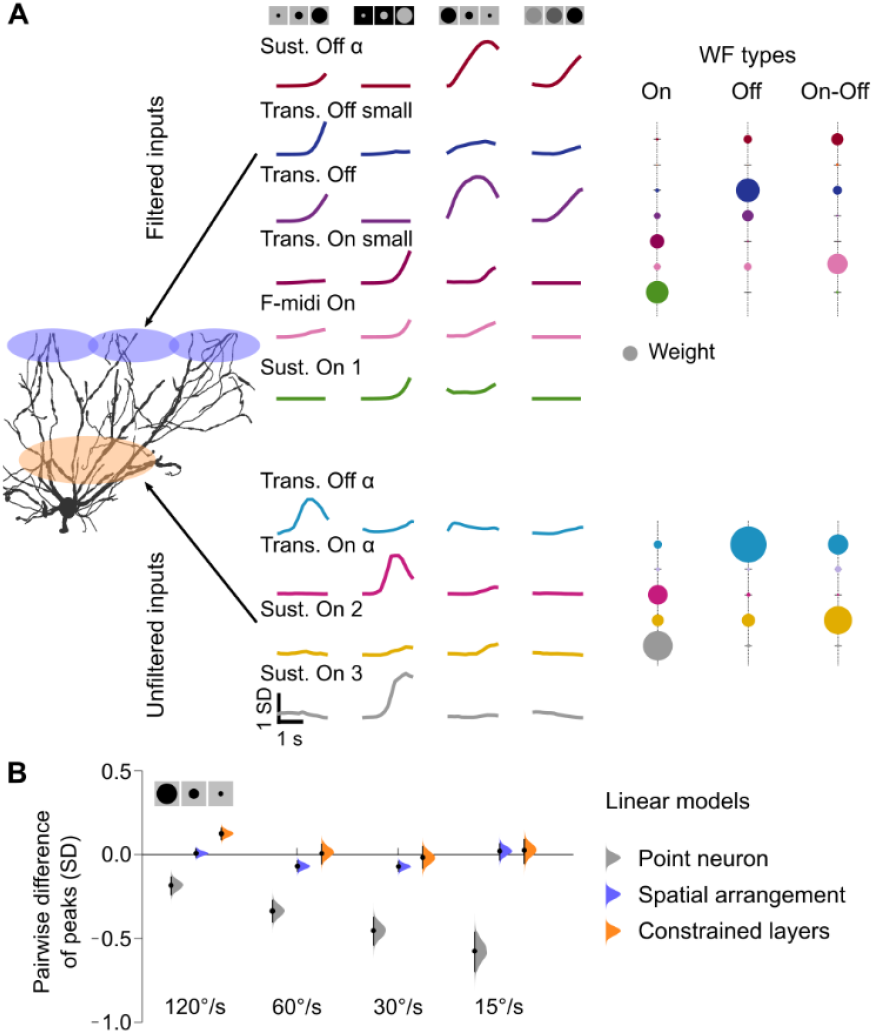
A Model with Layer-Specific Spatiotemporal Integration Captures Wide-Field Neuron Responses. (A) Model incorporating layer-specific spatiotemporal integration: Retinal inputs to upper layers (depth <150 µm; see Figure 2H) are filtered by a spatiotemporal kernel, while inputs to lower layers (depth >150 µm) remain unaltered. Left: Filtered inputs. Right: Weights of filtered inputs. (B) Pairwise differences between peak wide-field neuron responses and model fits, comparing models with spatially arranged inputs with and without constrained layers.

## Discussion

Our findings demonstrate that dendritic morphology and the detailed spatial organization of inputs are not merely a structural feature of the colliculus but a computational substrate that enables selective filtering of salient motion cues. Salient visual motion refers to movement patterns that stand out from the background and are likely to trigger an innate behavioral response. The layered organization of inputs along the extensive dendritic tree of wide-field neurons allows functionally distinct retinal signals to be processed in a spatially structured manner, supporting efficient multiplexing of different types of motion that mimic approaching, receding and passing objects. By selectively amplifying slow-moving edges, while filtering out static and fast-moving signals, wide-field neurons contribute to a reliable detection of environmental threats and opportunities.

### Wide-field neuron responses are not explained by retinal relay

Wide-field neurons perform a *de novo* computation of motion selectivity, as their responses to shrinking stimuli cannot be accounted for by a simple weighted sum of retinal inputs. Unlike the retina, where expanding stimuli dominate, wide-field neurons exhibit balanced tuning to both expanding and shrinking objects. This transformation arises from the structured integration of diverse retinal inputs, rather than from local inhibitory interactions. These findings suggest that dendritic processing in the colliculus extracts behaviorally relevant motion features in a way not explicitly present in the retinal input.

### Contribution of the layered organization of inputs

Our findings demonstrate that the dendrites of wide-field neurons receive inputs in a depth-specific manner, with different retinal ganglion cell types targeting specific layers of the colliculus. This arrangement provides a physical substrate for filtering and integrating distinct motion cues. Our estimates of the position of retinal ganglion cell innervation align with known stratification depths of different ganglion cell types, e.g. Foxp2+ and Smi32+ ganglion cells^2,46–48^. In addition, our estimates allow us to localize the input position of ganglion cell types that are not labeled in a specific mouse line. This has potential computational consequences. For example, Transient On and Off alpha cells restricted to the deepest layers (150-250 µm) likely relay looming responses, while their smaller counterparts, Small Transient On and Off cells in the superficial layers integrate signals over a large field of view, enabling spatial integration of small moving objects. Sustained Off alpha cells, in the intermediate layers (150-100 µm), may contribute to spatial integration of larger moving objects. Strong responses to receding stimuli likely arise from interactions between Sustained Off alpha cells, which respond early to large moving disks, and Small Transient cells in distal dendrites, activated by small moving disks. Inputs from Sustained Off alpha cells closer to the cell body could then facilitate the propagation and integration of signals from distal dendrites. Future studies could test this hypothesis by targeted activation and measurements of dendritic compartments in acute slices.

### Model complexity and the role of spatial organization

Our modeling approach reveals that incorporating layer-specific inputs allows us to explain wide-field neuron responses with minimal complexity. Each retinal ganglion cell type targets a specific layer and is assigned a layer-specific filter, reducing the need for additional computational mechanisms. This principle mirrors findings from artificial neural networks trained on visual tasks, where models with local connectivity that reflect retinotopic organization require three orders of magnitude fewer parameters than model with all-to-all connectivity while achieving the same performance^49,50^. Such efficiency is critical in biological systems, where rapid and reliable reactions are necessary for survival. By leveraging spatially organized input patterns, wide-field neurons effectively extract motion cues essential for innate behaviors, illustrating how anatomical structure enhances computational efficiency in neural circuits.

### The role of low pass filtering and nonlinear integration

Our best-performing model relies on linear integration and the spatial arrangement of dendrites and inputs, but temporal filtering and nonlinear integration may also contribute. First, neurons with large somas and long dendrites often exhibit strong low pass filtering, e.g., dopamine neurons (~80 ms time constant)^51^ and teleost neurons (~100 ms)^52^. However, models incorporating temporal delays best fit the data at ~1 s, far exceeding natural observations and instead reflecting the duration of the presented stimuli. Second, while wide-field neurons exhibit active dendritic properties^39^, nonlinear integration does not significantly shape their responses. Advanced multilayer perceptron models with nonlinearities provided only marginal improvements, suggesting that complex nonlinear processing is unnecessary. Instead, as described by Gale and Murphy, small stimulations of distant dendritic branches within the broad receptive field reliably generate somatic spikes^39^, leading to effective linear summation of inputs across dendrites.

### Functional subtypes of wide-field neurons might originate from subtype-specific wiring

We identified three functional subtypes of wide-field neurons with distinct contrast preferences. Our modeling and correlation analysis suggest that each subtype receives input from a specific subset of the identified On, Off and On-Off retinal ganglion cell types. Genetic subtypes with unique molecular profiles^53–56^ may underlie this subtype-specific connectivity, giving rise to the observed contrast selectivity.

### Caveats of calcium imaging

This study relies on fluorescent calcium signals, which present two main limitations: low temporal resolution and limited bandwidth. Low temporal resolution restricts our analysis to rate coding, while limited bandwidth necessitates careful consideration of the dynamic range of the measured signals. We deem calcium imaging an appropriate method, considering that with the current approach we measure retinal inputs and wide-field neuron outputs separately and hence do not aim to infer effects of spike time correlations. To address the potential impact of brightness saturation in calcium signals, we performed dual patch-clamp recordings of spiking activity and calcium signals in retinal ganglion cells. Additionally, wide-field neurons show low baseline activity (~0 Hz) and moderate firing rates (up to 50 Hz) in response to motion^25,39^, keeping our signals within the calcium indicator’s linear range^57^. However, our detection of dendritic signals is limited to events that induce local calcium influx, excluding synapses generating primarily sodium spikes. This may explain why we did not detect inputs from each of the 12 retinal ganglion cell types providing input to wide-field neurons. Future studies could overcome this limitation using dendritic patch-clamp recordings or voltage indicators optimized for *in vivo* imaging.

### Modulatory inputs likely have a minimal impact on visual integration properties of wide-field neurons

This study focused on two types of inputs: excitatory inputs from the retina and inhibitory inputs from Gad2 neurons of the colliculus. However, wide-field neurons receive input from many other brain regions^43,58–60^, many of which are thought to have modulatory roles, adjusting the output of these neurons based on the animal’s internal state or other sensory inputs. Since we analyzed median responses to repeated stimulus presentations, we expect these modulatory effects to average out. Additionally, one of the primary sources of modulation, the primary visual cortex - known to modulate the gain of collicular neurons in response to expanding disks^58^ - was surgically removed in our experiments to expose the anterior part of the superior colliculus for calcium imaging.

### Wide-field neurons of the colliculus are a versatile tool to study single neuron computations

Wide-field neurons of the colliculus are a powerful tool for studying single neuron computations and saliency processing, as they reliably drive experimentally tractable innate orienting behaviors^24,25,29,42–44^. These neurons can be genetically targeted using the *Ntsr1-GN209-Cre* mouse line or retrogradely traced from the lateral posterior nucleus of the thalamus, enabling precise experimental control^40,41^. Their well-defined behavioral outputs provide a direct link between neural perturbations and behavior, offering a unique platform to investigate how disruptions in dendritic properties impact neuronal function and organism level outcomes.

### Saliency processing in the colliculus

The superior colliculus rapidly extracts behaviorally relevant motion cues, enabling fast orienting responses. Unlike the retinothalamic pathway, which relays features encoded by the retina to the cortex^18,61^, wide-field neurons of the collliculus specialize in detecting motion features that demand immediate action. Our results suggest that dendritic processing in wide-field neurons selectively amplifies slow-moving edges while filtering out static and fast-moving signals. This selectivity may be critical for behavioral decisions based on the speed of approaching threats or escaping prey. Future behavioral studies using chemogenetic suppression of wide-field neurons could directly test their role in speed-dependent decision-making.

## ACKNOWLEDGEMENTS

We thank Bram Nuttin and Pedro Gonçalves (NERF, Leuven, Belgium) for discussions on the modeling aspects and Xu Han (VIB-CBD, Leuven, Belgium) who implemented the band pass noise stimulus. Initial rabies aliquots (Rabies-CVS-deltaG-GCaMP6f and Rabies-CVS-deltaG-Chr2-YFP) as well as cell lines for virus amplification (Neuro2a-G and Neuro2a-EnvA) were provided by Andy Murray (Sainsbury Wellcome Center, London, UK). This work was supported by the FWO (G094616N to KF, G091719N to KF, 1205421N to NKK, 12S7917N/12S7920N to KR); the European Union’s Horizon 2020 research and innovation program under the Marie Sklodowska-Curie grant agreement no. 796102 to NKK and no. 665501 to KR.

## AUTHOR CONTRIBUTIONS

Conceptualization, N.K.K., V.B. and K.F.; experimental setup and development, N.K.K., C.L., J.Z. and K.R.; in vivo imaging, N.K.K.; retinal imaging, C.L.; sparse labeling, J.Z.; analysis, N.K.K. and C.L.; computational models, N.K.K. and N.B.; software, N.K.K. and K.R.; writing & editing, N.K.K. and K.F.

## DECLARATION OF GENERATIVE AI AND AI-ASSISTED TECHNOLOGIES

During the preparation of this work, the authors used ChatGPT to check for consistency and correct sentence structure. After using this tool, the authors reviewed and edited the content as needed and take full responsibility for the content of the publication.

## Methods & Materials

### RESOURCE TABLE

| REAGENT or RESOURCE | SOURCE | IDENTIFIER |
| --- | --- | --- |
| **Antibodies** |  |  |
| Chicken, polyclonal, anti-GFP (1:500) | Thermo Fisher Scientific | Cat# A-10262<br>RRID: AB_2534023 |
| Donkey, Alexa 488, anti-chicken (1:500) | Immuno-Jackson | Cat# 703-545-155<br>RRID: AB_2340375 |
| Goat, polyclonal, anti-ChAT (1:200) | Millipore | Cat# AB144P<br>RRID: AB_11214092 |
| Donkey, Alexa 633, anti-goat (1:500) | Thermo Fisher Scientific | Cat# A-21082<br>RRID: AB_10562400 |
| Mouse, monoclonal, anti-SMI32 (1:1000) | Biolend | Cat# 801701<br>RRID: AB_2564642 |
| Donkey, Cy3, anti-mouse (1:400) | Immuno-Jackson | Cat# 715-165-151<br>RRID: AB_2315777 |
| Donkey, Dylight 405, anti-mouse (1:500) | Thermo Fisher Scientific | Cat# A-48257<br>RRID: AB_2884884 |
| Donkey, Dylight 405, anti-mouse (1:500) | ImmunoJackson | Cat# 715-475-150<br>RRID: AB_2340839 |
| Rabbit, polyclonal, anti-CART (1:500) | Phoenix Pharmaceuticals | H-003-62<br>RRID: AB_2313614 |
| Donkey, DyLight 405, anti-rabbit (1:400) | Immuno-Jackson | Cat# 715-475-150<br>RRID: AB_2340839 |
| Donkey, Alexa 555, anti-rabbit (1:500) | Thermo Fisher Scientific | Cat# A-31570<br>RRID: AB_2313501 |
| Goat, polyclonal, anti-FOXP2 (1:500) | abcam | Cat# 1307<br>RRID: AB_1268914 |
| Donkey, Alexa 555, anti-goat (1:300) | abcam | Cat# ab150130<br>RRID: AB_2927775 |
| Rabbit, polyclonal, anti-RFP (1:1000) | Rockland | Cat# 600-401-379<br>RRID: AB_2209751 |
| DAPI (1:200) | Roche | Cat# 10276236001 |
| **Bacterial and virus strains** |  |  |
| AAV1-hSyn-Flex-GCaMP6s | Addgene | Cat# 100845-AAV1<br>RRID: Addgene_100845 |
| EnvA-RVΔG-GCaMP6f:<br>EnvA-coated rabies CVS-N2c-ΔG-GCaMP6f | This paper based on aliquots obtained from Laboratory of Andy Murray | N/A |
| HSV-Flex-TVA-G-mCherry:<br>HSV-hEF1α-LS1L-TVA950-T2A-n2c(G)-IRES-mCherry | R. Neve (MGH) | RN716 |
| HSV-Flex-mCherry:<br>HSV-hEF1α-LS1L-IRES-mCherry | R. Neve (MGH) | RN413 |
| BHK cells | Laboratory of Botond Roska/ Laboratory of Karl-Klaus Conzelmann | N/A |
| B7GG cells | Laboratory of Botond Roska/ Laboratory of Karl-Klaus Conzelmann | N/A |
| BHK-EnvA cells | Laboratory of Botond Roska/ Laboratory of Karl-Klaus Conzelmann | N/A |
| HEK293T-TVA cells | Laboratory of Botond Roska/ Laboratory of Karl-Klaus Conzelmann | N/A |

| **Chemicals, peptides, and recombinant proteins** |  |  |
| --- | --- | --- |
| Lipophilic tracers DiI, DiD, DiO | Thermo Fisher Scientific | Cat# D7776, D7757, D275 |
| 10x PBS | VWR | Cat# 437117K |
| 1x PBS | VWR | Cat# 444057Y |
| Histofix 4% | Roche | Cat# P087.5 |
| Normal Donkey Serum | Millipore | Cat# 30-100ML |
| 10% Bovine Albumin | Sigma | Cat# SRE0036-250ML |
| DMEM, high glucose | Thermo Fisher Scientific | Cat# 41965062 |
| Trypsin 0.05% | Thermo Fisher Scientific | Cat# 25300054 |
| Fetal Bovine Serum (FBS) | Thermo Fisher Scientific | Cat# 10270106 |
| 2,2'-thiodiethanol (TDE) | Sigma | Cat# 166782-500G |
| ProLong® Gold Antifade Mounting Medium | Thermo Fisher Scientific | Cat# P36934 |
| Sodium Azide (NaN<sub>3</sub>) | Sigma | Cat# S2002-100G |
| Triton X-100 | Sigma | Cat# S8875 |
| Sodium Chloride (NaCl) | Sigma | Cat# S7653-250G |
| Potassium Chloride (KCl) | Sigma | Cat# P5405-25G |
| Calcium Chloride (CaCl<sub>2</sub>) | Sigma | Cat# C5670-100G |
| Magnesium Chloride (MgCl<sub>2</sub>) | Sigma | Cat# 4880 |
| D-glucose (Dextrose) | Sigma | Cat# D9434-250G |
| Sodium phosphate monobasic (NaH<sub>2</sub>PO<sub>4</sub>) | Sigma | Cat# S5011 |
| Sodium Hydroxide (NaOH) | Sigma | Cat# 655104-500G |
| Sodium bicarbonate (NaHCO<sub>3</sub>) | Sigma | Cat# S8875-1KG |
| **Deposited data** |  |  |
| Preprocessed raw data | This paper; Zenodo |  |
| **Experimental models: Organisms/strains** |  |  |
| Mus musculus Ntsr1-GN209-Cre: STOCK Tg(Ntsr1-Cre) GN209Gsat/Mmucd | Laboratory of Keisuke Yonehara | Genset: 030780-UCD |
| Mus musculus Gad2-IRES-Cre: STOCK Gad2tm2(cre)Zjh/J | The Jackson Laboratory | Cat# 010802<br>RRID: IMSR_JAX:010802 |
| **Software and algorithms** |  |  |
| Fiji | Schindelin et al. 2012 | RRID: SCR_002285 |
| MATLAB | Mathworks | RRID: SCR_001622 |
| Zen lite | Zeiss | N/A |
| CAFFE |  | caffe.berkeleyvision.org |
| ChAT band detector | This paper | https://github.com/farrowlab/ChATbandsDetection |
| VNET |  | github.com/faustomilletari/VNet |
| PYTHON 3.9 | Python Software Foundation | RRID: SCR_008394 |
| CaImAn 1.7.1 | Flatiron Institute | RRID: SCR_021533 |
| Vaa3D | Peng et al. 2014 | https://home.penglab.com/project/vaa3d |
| CANDLE | Coupe et al. 2012 |  |
| GNU Octave | Free Software Foundation | RRID: SCR_01439 |
| Psychophysics Toolbox | Psychtoolbox | RRID: SCR_002881 |
| WaveSurfer (version: 0.918) | Janelia Research Campus | RRID: SCR_021529 |
| ScanImage | Vidrio Technologies | http://scanimage.vidriotechnologies.com |
| **Other** |  |  |
| Rapid Flow Filters 0.2 μm pore size | VWR | Cat# 514-0027 |
| Wiretrol® II capillary micropipettes | Drumond Scientific | Cat# 5-000-2005 |
| Borosilicate glass | Sutter Instrument | Cat# BF100-20-10 |
| Glass capillaries, ∅ 3 mm, custom fine saw cut to L=1.5 mm | Hilgenberg | 1915017 |
| Round borosilicate cover glasses, ∅ 3 mm, 0.1 mm thickness | Multichannel systems | 640720 |
| UV-curing optical adhesive | Norland | 61 |
| Metabond dental cement | Crown & Bridge | N/A |
| Super-Bond C&B kit for self-curing adhesive resin cement | Sun Medical | N/A |
| Laser-Based Micropipette Puller | Sutter Instrument | Cat# P-2000 |
| Small Animal Stereotaxic Workstation | Narishige | Cat# SR-5N |
| Stereotaxic Micromanipulator | Narishige | Cat# SM-15R |
| Hydraulic Oil Micromanipulator | Narishige | Cat# MO-10 |
| Oil Microinjector | Narishige | Cat# IM-9B |
| Two-photon microscope | Scientifica | Serial# 14200 |
| 780 nm LED light source | Thorlabs | Cat# M780L3 |
| Patch-Clamp amplifier | Molecular Device | Axon Multiclamp 700B |
| Patch-Clamp microscope | Scientifica | Slice Scope |
| Patch-Clamp manipulator | Scientifica | Serial# 301311 |
| Zeiss LSM 710 confocal microscope | Zeiss | Cat# LSM710 |
| Mai Tai HP Ti:Sapphire laser | Spectra-Physics | N/A |
| B-Scope 2p microscope | Thorlabs | N/A |

## RESOURCE AVAILABILITY

### Data and code availability

Original code and analyzed data has been deposited at Zenodo and is publicly available as of the date of publication. The DOI is listed in the key resources table. Any additional information required to reanalyze the data reported in this paper is available from the lead contact upon request.

## EXPERIMENTAL MODEL AND STUDY PARTICIPANT DETAILS

### Mice

All experimental procedures were approved by the Ethical Committee for Animal Experimentation (ECD) of the KU Leuven and followed the European Communities Guidelines on the Care and Use of Laboratory Animals (014-2018/EEC, 166–2018/EEC). Both male and female adult transgenic mice (1–3 months old) were used in our experiments. To target wide-field neurons and their retinal inputs, *Ntsr1-GN209-Cre* mice were used (10 female, 5 male and 10 female, 5 male, respectively). Inhibitory neurons of the colliculus were targeted using *Gad2-IRES-Cre* mice (4 female, 4 male). Mice were heterozygously bread with C57Bl6J mice and kept on a 12-h light-dark cycle (lights on at 7:00), and sterilized food pellets and water were provided *ad libitum*.

## METHOD DETAILS

### Labeling of wide-field neurons and Gad2 neurons of the colliculus for two-photon imaging

To label wide-field neurons and Gad2 neurons of the superior colliculus for in vivo two-photon calcium imaging, a floxed viral vector expressing the fluorescent calcium indicator GCaMP6s (AAV1-hSyn-Flex-GCaMP6s, titer 2.956 × 10^13^ GC/mL, Addgene) is injected into the superior colliculus of *Ntsr1-GN209-Cre* and *Gad2-IRES-Cre* mice, respectively, and a cranial head plate and cannula window is implanted above the frontal two thirds of the superior colliculus, following a procedure previously developed in the lab^62^.

Anesthesia is induced at the beginning with Isoflurane 3% in 0.8 l/min O_2_ in a closed chamber and then kept with a combination of Ketamine (75mg/kg) and Medetomidine (1mg/kg). During the whole procedure animals are kept at physiological temperature by a homeothermic blanket. After deep anesthesia is achieved, eye ointment is applied to protect the eyes from drying out. Before starting any surgical procedure, the paw of the animal is pinched to check for the absence of pedal reflex (indicator of a proper anesthesia). The scalp is shaved and depilated (depilatory cream), disinfected (70% EtOH and betadine®), 0.5% Lidocaine is applied locally and the skin opened/removed; the periosteum is removed, and the lateral and posterior muscles are retracted. Vetbond TM is applied to open skin and exposed muscle. A head plate is attached to the skull using cyanoacrylate glue and Metabond®; a 3.5 mm craniotomy is performed; the brain is washed with sterile aCSF; a small cortex trunk (3 mm in diameter, 1 mm deep) is gently aspired under visual guidance until the surface of the SC is exposed. A cannula window (3 mm diameter, 3 mm height) is wedged into the craniotomy. Animals recover in separate cages and are treated with analgesic and antibiotics for 72 hours (Vetergesic, Ceva Animal Health Ltd; Emdotrim, ecuphar, BE-V235523) and monitored for 4 days.

### Labeling of retinal ganglion cells innervating wide-field neurons

Retinal ganglion cells that project to wide-field neurons were labeled with a transsynaptic viral tracing strategy, combining a floxed helper virus (HSV-hEF1α-LS1L-TVA950-T2A-n2c(G)-IRES-mCherry) with an EnvA-coated CVS-N2c-ΔG-GCaMP6f rabies virus in *Ntsr1-GN209-Cre* mice^41^. Animals were quickly anesthetized with Isoflurane (Iso-vet 1000mg/ml) and then injected with a mixture of Ketamine and Medetomidine (0.75 mL Ketamine (100 mg/mL) + 1 mL Medetomidine (1 mg/mL) + 8.2 mL Saline). Mice were placed in a stereotaxic workstation (Narishige, SR-5N). Dura tear (NOVARTIS, 288/28062-7) was applied to protect the eyes.

First, the helper virus was injected into the lateral pulvinar using micropipettes (Wiretrol® II capillary micropipettes, Drumond Scientific, 5-000-2005) with an open tip of around 30 µm and an oil-based hydraulic micromanipulator MO-10 (Narishige) for stereotactic injections. The injection coordinates for a 4 weeks old mouse with a bregma-lambda distance of 4.7 mm were AP: −1.85; ML: ±1.50; DV: 2.50 mm. As the mice were different in body size, we adjusted the coordinates for each mouse according to their bregma-lambda distance. To label the injection sites, DiD (Thermo, D7757) was used to coat the pipette tip. We injected in total 100-400 nl helper virus in single doses of up to 200 nl with a waiting time of 5-10 min after each injection.

Following the injections, the wound was closed using Vetbond tissue adhesive (3M,1469). After surgery, mice were allowed to recover on top of a heating pad and were provided with soft food and water containing antibiotics (emdotrim, ecuphar, BE-V235523).

Twenty-one days later, we injected the rabies virus into the superior colliculus using the same method as for the helper virus injections. As the retinotopic location of the first injection into the lateral pulvinar is uncertain, we covered as much as possible of the superficial layer of the superior colliculus during the second injection to maximize the labeling of cells in the retina. We injected 100-200 nl of rabies virus at a depth of 1.7 – 1.8 mm at the corners of a 1 mm^2^ square anterior of lambda and starting at the midline.

### Sparse labeling of wide-field neurons

We sparsely labeled wide-field neurons by injecting a diluted HSV-hEF1α-LS1L-mCherry (1:128) into the lateral posterior nucleus of the thalamus of *Ntsr1-GN209-Cre* mice (see Labeling of retinal ganglion cells).

### *In vivo* colliculus recordings

Before recordings started, animals were acclimated to the experimenter for a week. Mice were handled for 10 min a day for a period of three days. Mice were head fixed on an air-cushioned Styrofoam ball for progressive periods of time through five days, starting from 10 min to an hour.

The superior colliculus was scanned with a SpectraPhysics MaiTai femtosecond laser at 920 nm through a Thorlabs B-Scope microscope equipped with a resonance scanner and a Nikon 16x objective with 3 mm focal length. The laser power was adjusted and blanked during fly back of the galvo mirror by a Pockels cell. For imaging the somas of wide-field neurons, the nominal power at the end of the objective is kept at 90 mW while an area of 250 × 250 μm^2^ is imaged with a resolution of 512 × 512 pxls. This results in an imaging rate of 30 Hz. For imaging the dendrites of wide-field neurons, the superior colliculus is scanned across 5 planes with a piezo element starting at the surface and going down to 250 μm. The laser power is adjusted by depth following an exponential, starting at 30 mW and saturating at 120 mW. Fluorescence of the sample is filtered by a Thorlabs GFP emission filter (MF525-39) and photons captured by a Hamamatsu PMT.

The B-Scope is operated and imaging frames are recorded using ScanImage 11 (Vidrio Technologies). Imaging frames and visual stimuli are synchronized by yoking ScanImage to WaveSurfer 0.913 (HHMI Janelia Research Campus), which lets us capture imaging frame and visual stimulus triggers and the spinning speed of the Styrofoam ball. Further, square pulse triggers of 60 Hz are sent to a face camera (Mako G-030B, Allied Vision Technologies), recording the ipsilateral side of the mouse’s face. The camera frames are recorded with custom-written software by Joao Couto (bitbucket.org/jpcouto/labcams) in Python 2.7.

During calcium imaging, visual stimuli were displayed on a 78 × 40 cm^2^ LCD screen at 30 cm distance from the mouse and centered at 0° elevation and 20° azimuth of the contralateral eye. Visual stimuli were generated using a custom-written software in GNU Octave (Eaton et al. 2019) with Psychtoolbox-3.

### *Ex vivo* retina recordings

Visual responses of labeled retinal ganglion cells were recorded with two-photon calcium imaging (n = 1471 cells, 14 mice) or whole-cell loose patch recordings (n = 112 cells, 11 mice). Calcium responses were confirmed with simultaneous whole-cell patch recordings (n = 5 cells, 2 mice, Figure S4).

For *ex vivo* recordings of retinal ganglion cells, retinas were isolated from mice that were dark-adapted for a minimum of 30 minutes. Retina isolation was done under deep red illumination in Ringer’s medium (110 mM NaCl, 2.5 mM KCl, 1 mM CaCl2, 1.6 mM MgCl2, 10 mM D-glucose, 22 mM NaHCO3, bubbled with 5% CO2/95% O2, pH 7.4). The retinas were then mounted ganglion cell-side up on filter paper (Millipore, HAWP01300) that had a 3.5 mm wide rectangular aperture in the center, and superfused with Ringer’s medium at 32–36°C in the microscope chamber for the duration of the experiment.

Electrophysiological recordings were made using an Axon Multiclamp 700B amplifier (Molecular Devices) and borosilicate glass electrodes (BF100-50-10, Sutter Instrument). Signals were digitized at 20 kHz (National Instruments) and acquired using WaveSurfer software (version: 0.918) written in MATLAB. The spiking responses were recorded using the patch clamp technique in loose cell-attached mode with electrodes pulled to 3-5 MΩ resistance and filled with Ringer’s medium. To visualize the pipette, Alexa 555 was added to the Ringer’s medium.

Fluorescent cells were targeted for recording using a two-photon microscope (Scientifica) equipped with a Mai Tai HP two-photon laser (Spectra Physics) integrated into the electrophysiological setup. To facilitate targeting, two-photon fluorescent images were overlaid with the IR image acquired through a CCD camera. Infrared light was produced using the light from an LED. For some cells, z-stacks were acquired using ScanImage (Vidrio Technologies).

Retinas are visually stimulated with a LED projector whose light is passed through a lens system and dimmed by a neutral density filter at the end of the optical pathway. The retina is oriented in a way that the nasal direction is facing down with respect to the projected image. During two-photon imaging, the stimulus is presented in blue to avoid overstimulating the PMTs

Stimuli were generated with an LCD projector (Samsung, SP F10M) at a refresh rate of 60 Hz, controlled with custom software written in Octave based on Psychtoolbox. The projector produced a light spectrum that ranged from ~430 nm to ~670 nm. The power produced by the projector was 240 mW/cm2 at the retina. A combination of a neutral density filter and a short pass filter (cutoff wavelength = 475 nm) were used to control the stimulus intensity in logarithmic steps. Recordings were performed with filters decreasing the stimulus intensity by 1-2 log units.

### Visual stimuli

The following visual stimuli were presented to the explanted retina and the behaving mice (white, gray and black correspond to 100%, 0% and −100% Weber contrast). A TTL pulse was sent at the beginning and the end of each repeat for later alignment.

#### Full-field stimulus

A fixed sequence of full-field contrast modulation was presented across the full screen (90° x 50°). The sequence starts with 2 s black, 3 s white, 3 s black and 2 s gray screen and is followed by two sinusoidal temporal contrast modulations, interleaved by 2 s of gray screen. The first sinusoidal is of 100% contrast and increases linearly in frequency from 0 to 8 Hz within 8 s. The second sinusoidal is of 2 Hz frequency and linearly increases in amplitude from 0 to 100% within 8 s. The second sinusoidal is followed by 2 s gray and 2 s black screen^18,26^. The whole sequence is repeated 10 times.

#### Expanding disk

A black or white disk linearly expanded from 2° to 50° within 0.95 s at the center of a gray or black screen, respectively. The stimulus was repeated 10 times, interleaved by 3 s of gray screen.

#### Shrinking disk

A black disk linearly shrunk from 50° to 2° of diameter within 0.95 s at the center of a gray screen. The stimulus was repeated 10 times, interleaved by 3 s of gray screen.

#### Dimming disk

A disk of 50° diameter linearly dimmed from background gray to black within 0.95 s at the center of the screen. The stimulus was repeated 10 times, interleaved by 3 s of gray screen.

#### Sweeping disk

A black or white disk of different diameter (2°, 8° and 32°) moved with 30°/s and 160°/s in two directions (left to right, right to left) across the center of a gray or black screen, respectively. Each combination of direction, size and speed was repeated 3 times in random order and interleaved by 3 s of gray screen.

#### Bandpass noise (in vivo only)

A patch of 50° was shown for 2 s on gray background with band-pass-filtered spatiotemporal white-noise patterns of different temporal (0.5, 1, 2, 4, 8 Hz) and spatial cut-off frequencies (0.02, 0.04, 0.08, 0.16, 0.32 cycles/°). Each pattern was randomly generated, and each condition repeated 4 times in random order, interleaved by 3 s of gray screen.

### Retina immunohistochemistry

After *ex vivo* recordings, retinas were fixed in 4% paraformaldehyde (Histofix, ROTH, P087.5mm) with 100 mM sucrose for 30 min at 4°C, and then transferred to a 24-well plate filled with 1x PBS and washed 3 times for 10 min at room temperature or transferred into 15 ml 1x PBS and washed overnight or longer at 4 °C. After washing, retinas were transferred to wells containing 10% sucrose in 1x PBS with 0.1% NaN3 (w/v) and allowed to sink for a minimum of 30 min at room temperature. Then retinas were transferred to wells containing 20% sucrose in 1x PBS with 0.1% NaN3 (w/v) and allowed to sink for a minimum of 1 hour at room temperature. Finally, retinas were put into 30% sucrose in 1x PBS with 0.1% NaN3 (w/v) and allowed to sink overnight at 4°C. The next day, freeze-cracking was performed: retinas were frozen on a slide fully covered with 30% sucrose for 3-5 min on dry ice. The slides were then thawed at room temperature. The freeze-thaw cycle was repeated two times. Retinas were washed 3 times for 10 min each in 1x PBS, followed by incubation with blocking buffer (10% NDS, 1% BSA, 0.5% TritonX-100, 0.02% NaN3 in 1x PBS) for at least 1 hour at room temperature or overnight at 4°C with gentle shaking.

Primary antibody goat against FOXP2 (abcam1307, 1:2000) was added after blocking and retinas were incubated for 5 days under constant gentle shaking at 4°C. They were prepared in 5% NDS, 0.3% TritonX-100 in 1x PBS. After incubation, retinas were washed 3 times for 15 min in 1x PBS with 0.3% TritonX-100 before being transferred into the secondary antibody solution (Alexa555 donkey anti-goat abcam150130, 1:300); prepared in 1xPBS overnight at 4°C. The second day, retinas were washed 3 times in 1x PBS and incubated in the second primary antibody solution for 5 days under constant gentle shaking at room temperature. The second primary antibodies were rabbit anti-GFP (Invitrogen, A-11122, 1:500), goat anti-ChAT (Chemicon, Ab144P, 1:200) and mouse SMI32 (Biolend, 801701,1:1000). They were prepared in 3% NDS, 1% BSA, 0.5% TritonX-100, 0.02% NaN3 in 1x PBS. Retinas were then washed 3 times for 10 min in 1x PBS with 0.5% TritonX-100. After washing, the retinas were incubated in the secondary antibody solution overnight at 4°C. Secondary antibodies were Alexa488 donkey anti-chicken (ImmunoJackson, 703-545-155, 1:500), Alexa 633 donkey anti-goat (Invitrogen A-21082, 1:500) and DyLight 405 donkey anti-mouse (Thermo, A-48257 or ImmunoJackson, 715-475-150, 1:300); prepared in 3% NDS, 1% BSA, 0.5% TritonX-100, 0.02% NaN3 in 1x PBS). Retinas were then washed 3 times in 1x PBS with 0.5% TritonX-100 and 1 time in 1x PBS.

For mounting, we used 2,2′-Thiodiethanol (TDE) (Sigma, 166782-500G) (Staudt et al., 2007) to exchange the water in the sample. To achieve this, retinas were incubated in different concentration of TDE buffer (10% −> 25% −> 50% −> 97%) for at least 30 minutes each. Then the retinas were embedded in ProLong® Gold Antifade Mountant (Thermo, P36934) and gently covered with a #0 coverslip (MARIENFEL, 0100032, No.0, 18*18 mm). To avoid squeezing the retinas, we put 4 strips of Parafilm (Parafilm, PM999) around the retina before adding the coverslip. Some of the retinas were mounted in 97% TDE with DABCO (Sigma, 290734) after immersion into TDE. Some retinas were mounted with ProLong® Gold Antifade Mountant directly after washing. Afterward, nail polish was used to prevent evaporation and the samples were stored in darkness at 4°C.

### Brain immunohistochemistry

Extracted brains were post-fixed in 4% PFA overnight at 4°C. Vibratome coronal sections (100 mm) were collected in 1x PBS and were incubated in blocking buffer (1x PBS, 0.3% Triton X-100, 10% Donkey serum) at room temperature for at least 1 hour or overnight. Then brain slices were incubated with primary antibodies solution for 2-3 days at 4°C with shaking. Slices were later washed 3 times for 10 min each in 1x PBS with 0.3% TritonX-100 and incubated in secondary antibody solution for 2-3 days at 4°C. Primary and secondary antibody were chicken anti-GFP (1:1000) and Alexa488 donkey anti-chicken (1:800), respectively, for GCaMP6s labeled cells and rabbit anti-RFP (1:1000) and Alexa555 donkey anti-rabbit (1:500) for sparsely labeled cells. Nuclei were stained with DAPI (1:1000) together with the secondary antibody solution. Sections were then again washed 3 times for 10 min in 1x PBS with 0.3% TritonX-100 and 1 time in 1x PBS, covered with mounting medium and a glass coverslip. PBS were prepared with 0.02% NaN3.

### Confocal microscopy

Confocal microscopy was performed on a Zeiss LSM 710 microscope. Overview images of the retina and brain were obtained with a 10x (plan-APOCHROMAT 0.45 NA, Zeiss) objective. The following settings were used: zoom 1.2, 5×5-tiles with 15% overlap, 2.37 µm/pixel resolution. For single retina ganglion cells and sparsely labeled wide-field neurons, we used a 63x (plan-APOCHROMAT 1.4 NA, Zeiss) objective. The following settings were used: zoom 0.7, 2×2-tiles or more (depending on size and number of cells) with 15% overlap. This resulted in an XY-resolution of 0.38 µm/pixel and a Z-resolution between 0.3 µm/pixel. The Z-stacks covered approximately 50 µm in depth.

## QUANTIFICATION AND STATISTICAL ANALYSIS

Data processing and analysis was performed in Python and MATLAB.

### Two-photon image registration and ROI extraction

Two-photon imaging frames were registered, and ROIs determined using NNMF with CaImAn 1.7.1 (Flatiron Institute) in Python 3.4. The signal decay time for the model was adjusted to GCaMP6s (1.4 s). ΔF/F values of these selected ROIs were used for further analysis. ROIs of collicular wide-field neurons, Gad2 neurons and retinal ganglion cells were obtained separately from ROIs representing dendritic compartments of collicular wide-field neurons.

### Clustering of ROI responses

Calcium responses were baseline-subtracted by the minimum and normalized by the standard deviation across all stimuli. K-Means clustering was performed on the correlation matrix of concatenated responses. For wide-field neurons, responses to expanding, shrinking, and dimming stimuli were used. Dendritic compartments were clustered based on full-field responses, while Gad2 neurons and retinal ganglion cells were clustered using responses to full-field, expanding, shrinking, and dimming stimuli.

### Cell selection

The repeats of visual stimuli and neural responses were aligned by using the simultaneously recorded stimulus start and imaging frame triggers. For each presented stimulus, a quality index of each ROI was calculated. The quality index was calculated from the variance of ΔF/F across time, var(·)_t_, and the mean across stimulus repeats, E(·)_r_, as follows: QI = var(E(ΔF/F)_r_)_t_ / E(var(ΔF/F)_t_)_r_. Cells were included into the analysis if they were visually responsive with a quality index above 0.6.

### Identification of retinal ganglion cell types

We identified the recorded retinal ganglion cells in the stained retinas by template matching (FIJI plugin), allowing us to link visual responses to molecular types. For a subset (n = 122 cells, 12 retinas), the dendritic arbors were traced, allowing for morphological classification. Using the *On*- and *Off* ChAT bands, formed by the starburst amacrine cells, as landmarks of retinal organization, we calculated the stratification density profile of every traced retinal ganglion cell and measured the soma size. Retinal ganglion cells were then identified based on molecular labeling, soma size and stratification profile by matching them manually to existing databases^26–28,41^.

### Morphology of retinal ganglion cells

To annotate dendritic trees in confocal Z-stacks, we used Ariadne-service GmbH (Switzerland) for automated tracing. Prior, the confocal Z-stacks of retinal ganglion cells were denoised (CANDLE package^63^, MATLAB) and down-sampled (XYZ = 0.5 × 0.5 × 0.35 µm per pixel). ChAT band positions were extracted using a convolutional neural network (V-Net) trained to detect and segment these structures (https://github.com/farrowlab/ChATbandsDetection). The ChAT band locations were then used to warp the dendritic tree, aligning it in 3D space^28^.

Dendritic tree depth profiles were computed by normalizing Z-positions and applying a low-pass Fourier filter. The dendritic area was estimated via convex hull approximation, with diameters calculated as D = 2*(area / π)1/2. The soma area was measured manually using the convex hull tool in ImageJ.

### Morphology of sparsely labeled wide-field neurons

A sparsely labeled wide-field neuron was traced and stitched across four coronal slices in Vaa3D. The extracted start and end points of dendritic segments and their thickness were used to plot a coronal projection of the neuron.

### Depth distribution of dendritic clusters and retinal ganglion cell inputs

we determined the number of ROIs per scanning plane and corrected for sampling bias by weighting them relative to the total ROIs in each plane, yielding a depth distribution of dendritic ROIs per cluster.

To estimate the stratification depth of each retinal ganglion cell type, we identified the wide-field neuron dendritic ROI with the strongest correlation coefficient (>0.4) for each retinal ganglion cell. The depths of these identified dendritic ROIs were then used to calculate a depth distribution for each retinal ganglion cell type using the method above.

### Sparsity index

To test the amount of convergence and divergence of retinal inputs onto different dendritic clusters, we calculated the Gini index G for each row and column of the weight matrix, respectively. The distributions were compared to a uniform distribution across clusters and cell types.

### Expansion selectivity index

An expansion selectivity index was calculated for each recorded wide-field neuron and retinal ganglion cell based on peak responses to dark expanding disks r_e_ and bright expanding, shrinking and dimming disks r_s_ using the formula I = (r_e_ – r_s_)/(r_e_ + r_s_). Distributions were then visualized using kernel density estimation, with a smoothing bandwidth determined by Scott’s rule.

### Computational models

Models were implemented in Julia 1.7.

### Non-negative linear regression

A non-negative linear regression model^64^ was fitted using the NNLS package in Julia. The nonnegative constraint is in line with the excitatory nature of the retinal inputs to the colliculus and ensures sparsity.

Nonnegative linear regression was used in four different models, 1) a point neuron model integrating inputs from retinal ganglion cell types, 2) a point neuron model integrating inputs from retinal ganglion cells types and inhibitory interneurons, 3) a model considering low pass filtering of the inputs and 4) a model considering the spatiotemporal integration of distributed inputs.

To fit the responses of wide-field neuron cell bodies to expanding, shrinking and dimming disks, we concatenated the responses to the presented stimuli and normalized the concatenated responses by their standard deviation.

### Multilayer Perceptron

A multilayer perceptron was fitted using the Flux package in Julia. This model assumes a local summation of the excitatory retinal and inhibitory collicular inputs and nonlinear processing along the dendrites. A sigmoidal nonlinearity is used to capture supra- and sublinear processing of the summed inputs along the dendrites. To implement the neural spiking threshold, a rectifying nonlinearity is used at the output node.

Models were fitted by gradient descent using the ADAM optimizer. An L1 penalty on the sum of all fitted parameters was introduced to ensure sparsity. Linear weights and biases were initialized with uniformly distributed random values between 0 and 1.

### Models incorporating low pass filtering of inputs through dendrites

The non-negative linear regression model (see above) was fitted to an extended set of filtered retinal inputs, doubling the number of fitting parameters. A 4th-order Butterworth low-pass filter (cutoff: 0.4, 0.5, 1, 2, 3 Hz) generated additional inputs from the original inputs, and non-negative weights were fitted to the combined set of original and filtered inputs for each cutoff frequency.

### Models incorporating the spatial organization of inputs and dendrites

To capture for the spatial arrangement of inputs and dendrites, a spatiotemporal activation kernel was convolved with retinal responses to expanding and shrinking disks. This kernel accounted for the relative number of activated retinal inputs per 0.13 s time bin based on changes in disk surface area, producing a quadratically increasing or decreasing activation profile. The non-negative linear regression model (see above) was then fitted to the combined set of original and convolution-filtered retinal inputs.

To account for the layer-specific arrival of inputs at the dendrites, we refined our model by depth-dependent filtering. Retinal inputs to the upper dendritic layers (<150 µm), identified through correlation with dendritic signals, underwent spatiotemporal filtering, while inputs to the lower layers (>150 µm) remained unaltered.

