## Supplemental Information for "Dendritic Architecture Enables de Novo Computation of Salient Motion in the Superior Colliculus"

### Supplemental Figures

#### Figure S1 – related to Figure 1.

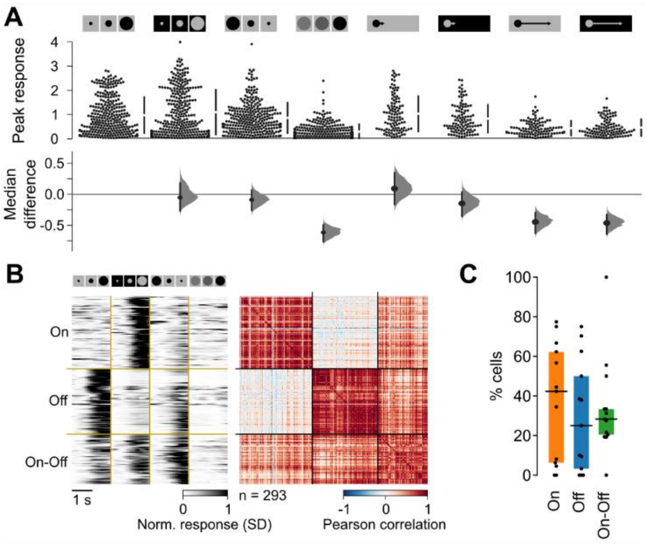


***Figure S1. Peak Responses and Clustering of Wide-Field Neurons***

*(A) Comparison of peak responses to expanding, shrinking and dimming disks and slow and fast 2° sweeping disks.*

*(B) Left: Heatmap normalized and clustered wide-field cell body responses. Right: Correlation matrix of response clusters.*

*(C) Proportion of On, Off, and On-Off wide-field cell bodies per recorded field of view. Vertical bars and boxes indicate median and interquartile range (IQR). Data as in Figure 1.*

**Figure S2 – related to Figure 1.**

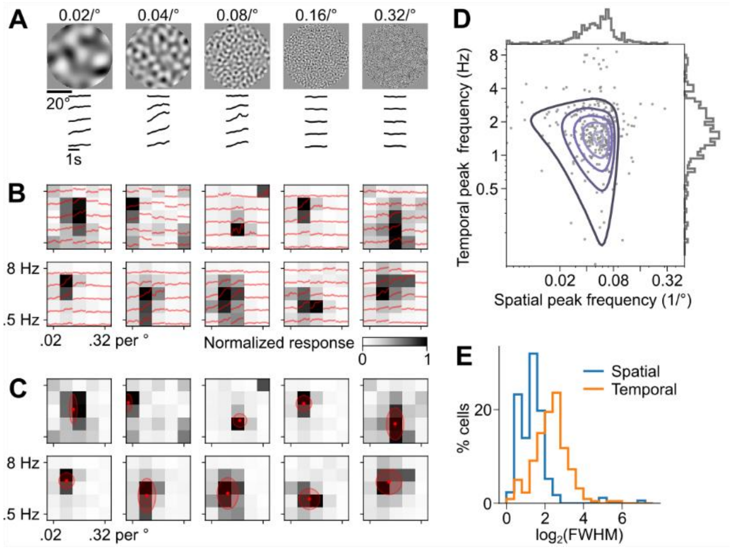


***Figure S2. Spatiotemporal Frequency Tuning of Wide-Field Neurons***

*(A) Responses of an example wide-field neurons to two second snippets of spatiotemporal bandpass noise of different spatial (left to right) and temporal frequencies (top to bottom).*

*(B) Spatiotemporal frequency tuning curves of ten example wide-field cell bodies, with heatmap indicating normalized peak responses.*

*(C) 2D Gaussian fits to the tuning curves of example cell bodies. Red dots indicate the mean, and shaded circles represent the 1-σ range.*

*(D-E) Distributions of the spatiotemporal peak frequencies (D) and tuning widths (E) of the 2D Gaussian fits. Data as in Figure 1.*

#### Figure S3 – related to Figure 1.

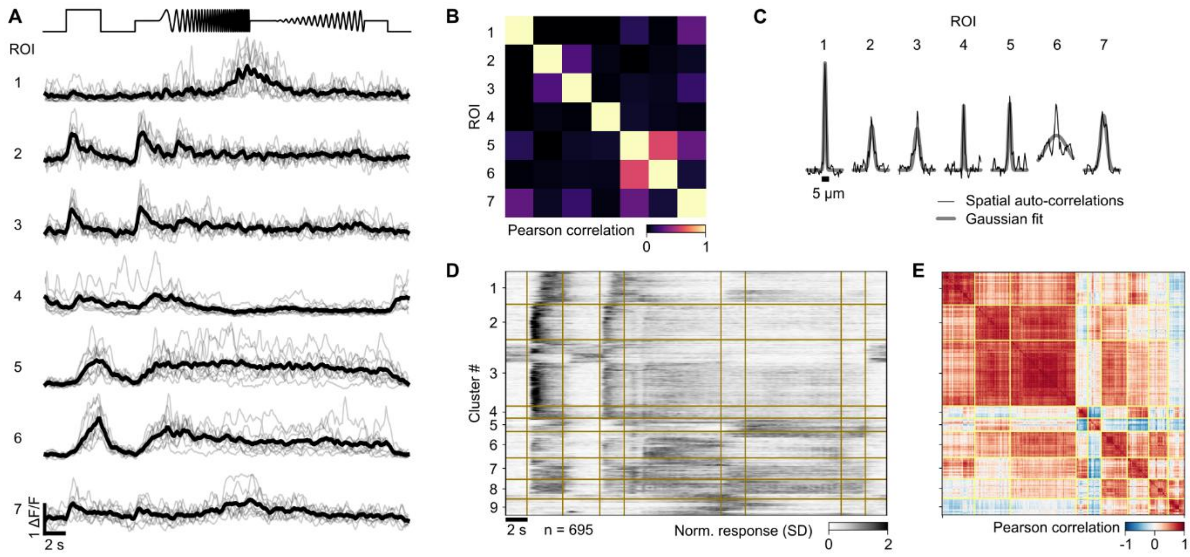


***Figure S3. Calcium Signals in Wide-Field Neuron Dendrites Are Highly Localized***

*(A) Example responses of dendritic ROIs to the full-field stimulus across four different imaging planes from a single recording. Gray and black lines indicate individual and averaged trials from 10 stimulus repetitions.*

*(B) Temporal correlations of raw ROI responses in (A) across all trials.*

*(C) Spatial auto-correlations of ROIs across the widest dimension.*

*(D-E) Heatmap of normalized and clustered average responses (D) and correlation matrix (E). Data as in Figure 1.*

**Figure S4 – related to Figure 2.**

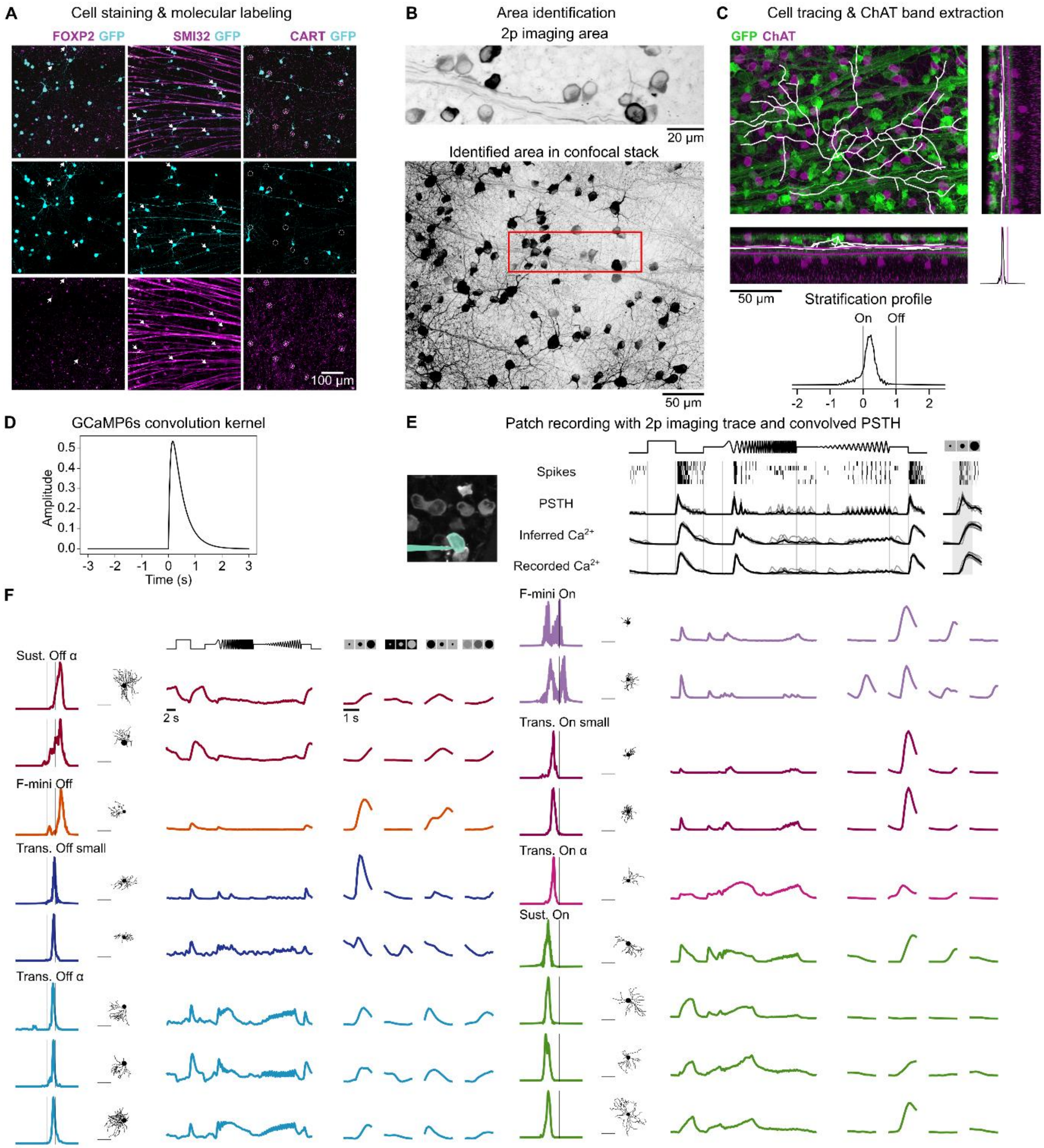


***Figure S4. Identification of Retinal Ganglion Cells Projecting to Wide-Field Neurons***

*(A) Immunohistochemical labeling of retinal ganglion cells. GFP-labeled cells (cyan) co-stained for FOXP2, SMI32, or CART (magenta) to identify molecular subtypes.*

*(B) Post hoc identification of the two-photon imaging area (top) within a stained whole-mount retina (bottom). Both images are maximum projections of z-stacks, with the imaging area (red box) determined via template matching.*

*(C) High-resolution confocal imaging of recorded cells (0.3765 × 0.3765 × 0.3 $\mu m^3$/pxl). Top: dendrite tracing of GFP-labeled cells (green). Bottom: extracted stratification profile relative to ON- and OFF-ChAT bands, with traced dendrites (white) and detected ChAT bands (purple).*

*(D) GCaMP6s convolution kernel used to convolve spike responses.*

*(E) Comparison of patch-clamp PSTHs with two-photon calcium signals from GCaMP6f and PSTHs convolved with the GCaMP6s kernel. Boxplots show correlations between retinal ganglion cell clusters and wide-field neuron clusters.*

*(F) Example retinal ganglion cells identified by molecular labeling, stratification profile, dendritic tree size, and visual responses. Data as in Figure 2.*

#### **Figure S5 – related to Figure 3.**

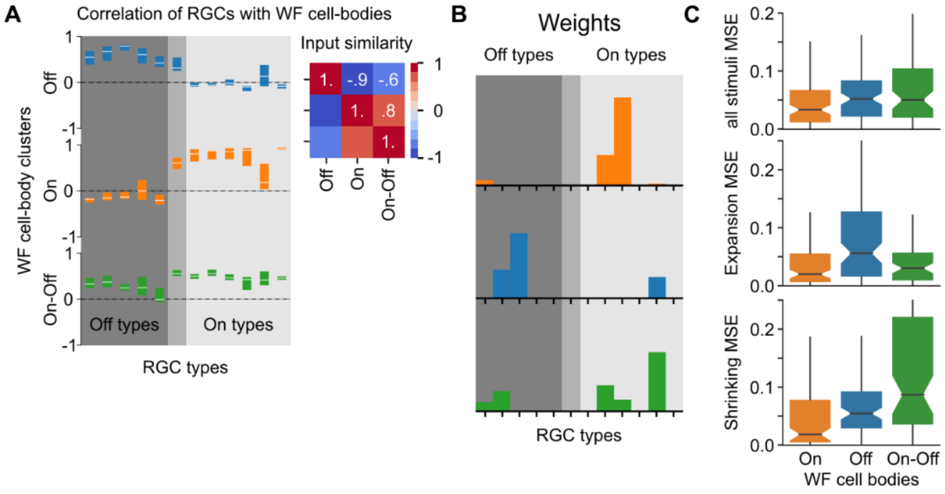


***Figure S5. Contribution of Retinal Ganglion Cells to Wide-Field Neuron Signaling***
*(A) Correlation between retinal ganglion cell types and wide-field neuron subtypes. Boxplots show median and interquartile range.*
*(B) Fitted weights for predicting average wide-field neuron subtype responses.*
*(C) Mean squared error (MSE) of response reconstruction across wide-field neuron clusters. Boxplots show median, interquartile range, confidence interval of the median (nudges), and whiskers extending to 1.5× the interquartile range.*

**Figure S6 – related to Figure 4.**

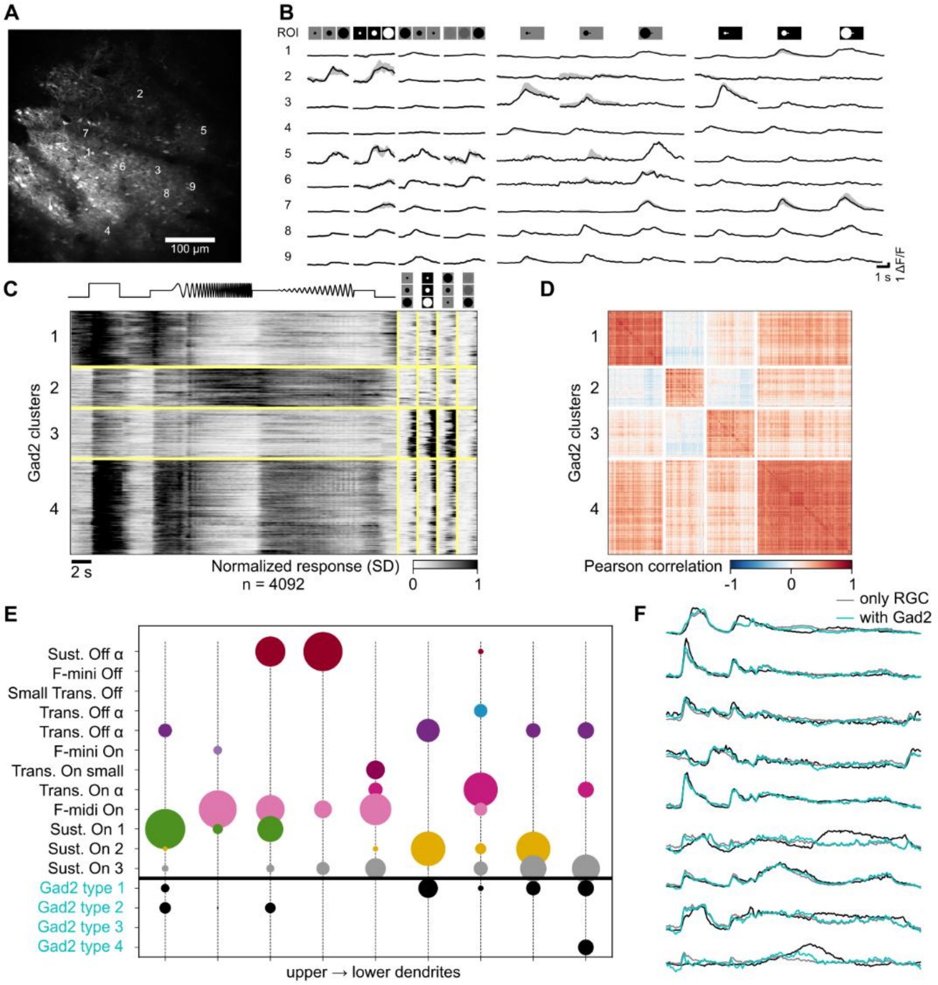


***Figure S6. Clustering of Gad2 Neurons and Contribution to Responses of Wide-Field Neuron Dendrites***

*(A) Sample recording of Gad2 neurons in the colliculus.*

*(B) Median and interquartile range of calcium responses of cells indicated in (A).*

*(C) Heatmap of normalized and clustered responses to full-field, expanding, shrinking and dimming stimuli.*

*(D) Correlation matrix of cell responses in (C).*

*(E) Weights of the non-negative linear fit of wide-field dendritic responses including Gad2 neurons.*

*(F) Linear fits of wide-field dendritic responses (black) with only retinal ganglion cells (gray) or including Gad2 neurons (cyan).*

**Figure S7 – related to Figure 5.**

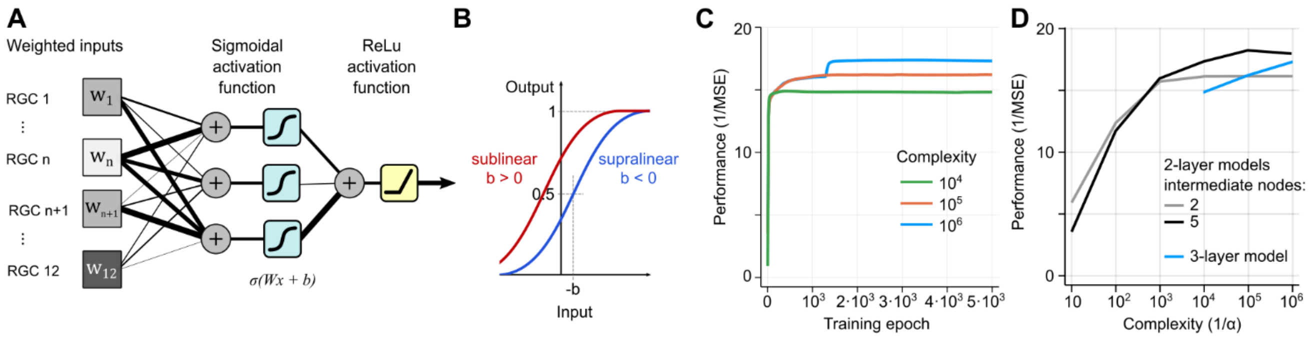


***Figure S7. Training of the Multilayer Perceptron and Model Performance by Complexity***

*(A) Architecture of the multilayer perceptron (MLP) model, consisting of two layers with three hidden nodes using sigmoidal activation functions and a single output node with a rectifying nonlinearity. The model pools inputs from twelve weighted retinal ganglion cell types.*

*(B) Sigmoidal nonlinearity with positive and negative bias terms (b), accounting for supralinear and sublinear integration of inputs, respectively.*

*(C) Model performance during training tested on withheld data. Model parameters (weights W and biases b) are regularized using an L1 cost function scaled by α, controlling the trade-off between model complexity and performance. Model complexity is inversely proportional to α (1/α), while performance is evaluated as the inverse of the mean squared error (1/MSE). Only models with high complexity outperform linear models.*

*(D) Model performance by complexity for different 2-layer and 3-layer models.*
